# Autumn drought drives deterministic bacterial filtering and network destabilization in a phenotype-related manner in *Pinus halepensis* seedlings

**DOI:** 10.64898/2026.05.11.724195

**Authors:** I Aleksieienko, IM Reiter, J Reilhan, M de Castro, C Santaella

**Affiliations:** Aix Marseille Univ, CEA, CNRS, BIAM, LEMiRE, ECCOREV FR 3098, F-13108 Saint-Paul-Lez-Durance, France; CNRS, Aix Marseille Univ, FR3098, ECCOREV, F-13545 Aix-en-Provence, France; ONF, PNRGF Cadarache, F-13108 Saint Paul Lez Durance, France

## Abstract

Mediterranean forest restoration depends critically on autumn seedling outplanting and establishment, a period increasingly threatened by delayed and irregular precipitation. Although critical, drought at moderate temperatures, decoupled from summer heat stress, remains poorly characterized in terms of plant physiology and root microbiome responses. In the present study we simulated a short but severe drought under moderate air temperatures (5–25°C) in *Pinus halepensis* Mill. seedlings to examine the independent effects of water deficit on physiology and root-associated microbial communities. Drought reduced stomatal conductance to one-third of control values and induced a decoupling between stomatal conductance and net photosynthesis upon rewatering. This decoupling is rather due to the residual hydraulic and biochemical limitations rather than transpirational cooling demands. Drought-treated seedlings diverged into distinct phenotypic classes differing in recovery capacity, with a subset failing to recover despite being phenotypically identical to the controls. Root microbiome restructuring was phenotype-dependent and differed between active and resident fractions: bacterial richness and evenness increased while bacterial assembly shifted progressively toward determinism with increasing phenotype severity (29% to 37%), whereas fungal communities shifted toward stochastic drift (up to 89%). Functionally, drought disrupted symbiotic associations and drove a shift toward a fungal saprotrophic lifestyle. Network analyses revealed displacement of *Rhizobium* from its central hub position, reducing connectivity and compromising functional resilience. These results demonstrate that short, severe drought at moderate temperatures fundamentally affect plant-microbiome interactions through phenotype-dependent assembly processes, with direct consequences for seedling establishment and Mediterranean reforestation under climate change.

## I. Introduction

Over recent decades, anthropogenic climate change has driven widespread losses of biodiversity and forest cover (Calvin et al., 2023), with particularly severe consequences in sub-arid Mediterranean regions (Lazoglou et al., 2024). In these ecosystems, reforestation practices rely on autumn or early winter planting historically timed to coincide with seasonal rainfall peaks that support root establishment before the onset of summer drought (Oliet et al., 2009). However, climate change is increasingly disrupting precipitation regimes, leading to unpredictable or delayed autumn rainfall and exposing newly planted seedlings to water deficits, a major driver of early mortality in Mediterranean reforestation programs (IGN, 2024; Pausas et al., 2004)

Despite this context, ecophysiological drought research has disproportionately focused on spring and summer conditions, reflecting a seasonal compartmentalization of stress studies and a prevailing emphasis on drought intensity rather than its timing (Charrier et al., 2021; Forner et al., 2018). This focus is partly rooted in the view that tree physiological activity declines outside the main growing period, leading to reduced consideration of off-season drought effects. Yet, this timing critically shapes tree physiological responses (Forner et al., 2018). This drought-heat framework is poorly suited to Mediterranean evergreen conifers such as *Pinus halepensis* Mill., a key species for reforestation in Southern Europe. Unlike species with a single growing season, *P. halepensis* exhibits a bimodal growth phenology with active cambial phases in both spring and autumn, separated by a summer drought-induced arrest (Camarero et al., 2010; Luis et al., 2007). Autumn therefore represents not an off-season but a second physiologically active window, during which soil moisture deficits constrain cambial growth, root establishment, and hydraulic function, particularly in newly transplanted seedlings where incomplete root–soil contact limits hydraulic recovery (Earles et al., 2018). Consistent with its thermophilic and predominantly isohydric strategy, *P. halepensis* relies on early stomatal closure and rapid hydraulic adjustment to avoid drought stress (Baquedano and Castillo, 2006; Moreno et al., 2024; Royo et al., 2001; Syropli and Iakovoglou, 2021), further increasing its sensitivity to water deficits during this critical establishment phase.

Seedling production is a cornerstone of reforestation programs, enabling the optimization of growth conditions and physiological status prior to out planting. Nursery practices allows fine-tuning of growing conditions, supporting root system development, physiological maturation, and the accumulation of carbohydrate reserves under optimal irrigation (Jaenicke, 1999; Lamhamedi et al., 2023; Sharpe, 1999). Prior to outplanting, seedlings can undergo a “hardening” phase that progressively acclimates them to field conditions through controlled water limitation (Finley et al., 2024; Puértolas et al., 2024). Despite this preparation, the transition from controlled conditions to the field often induces “transplant shock” (Mainhart et al., 2024; South et al., 2023), which can drive high seedling mortality when acclimation fails, particularly under water-limited conditions (Haase et al., 2021; Harrison et al., 2023).

Symbiotic and associative microorganisms play a crucial role in plant acclimation and drought resistance. Plant-associated microbiomes, comprising both resident and metabolically active fractions (Baldrian et al., 2012), enhance stress tolerance through phytohormone production, nutrient mobilization, and modulation of environmental perception (De Vries et al., 2020; Hoefle et al., 2024; Metze et al., 2023). Root exudation patterns shape microbiome composition and mediate plant-microorganism interactions (Berrios et al., 2023). In response to drought-induced plant stress signals, microbiome feedbacks modulate plant physiological responses, forming an integrated adaptive system at the intersection of plant physiology, root chemistry, and microbial ecology (Janse van Rensburg et al., 2024; Malik and Bouskill, 2022; Rolfe et al., 2019; Štraus et al., 2025).

While genotype-phenotype relationships in trees have been extensively characterized (Depardieu et al., 2021; Großkinsky et al., 2015; Howe et al., 2003; Mendez-Cea et al., 2023; O’Malley and Ecker, 2010), far less is known about the functional dynamics at the phenotype-microbiome interface. Understanding these dynamics is critical for improving plant stress tolerance and disease resistance in managed forest systems (Mendez-Cea et al., 2023; Trivedi et al., 2020), and for advancing ecosystem restoration through enhanced nutrient cycling and soil health (Compant et al., 2021; Vandenkoornhuyse et al., 2015). Most studies investigating microbiome responses in conifers have focused on summer drought or spring growing-season stress. This leaves the response of both plant and microbiome to autumn drought largely unexplored, particularly during the critical seedling establishment phase, characterized by limited local adaptation and high sensitivity to water (Candido-Ribeiro and Aitken, 2024).

To address these knowledge gaps, we conducted a controlled drought experiment on 7-month-old *P. halepensis* seedlings in autumn under greenhouse conditions, simulating off-season drought events increasingly observed in Mediterranean reforestation contexts. We aimed to determine how short-term, severe autumn drought affects seedling physiological response, including stomatal conductance, net photosynthesis, and growth and phenotypic traits, based on drought symptom severity and recovery capacity, and how these changes are associated with shifts in the composition and potential functionality of bacterial and fungal communities. We tested whether drought (i) alters the taxonomic and functional profiles of both resident and metabolically active microbiomes in a phenotype-dependent manner, (ii) modifies microbial assembly processes differentially across kingdoms and drought phenotypes, and (iii) restructures bacteria–fungi interaction networks, restructures bacteria–fungi interaction networks in ways that parallel host physiological state, thereby contributing to seedling phenotypic plasticity.

## II. Materials and methods

### 1. Plant material and growing conditions

Aleppo pine (*Pinus halepensis* Mill.) seedlings were obtained from seeds (origin: Mediterranean region) according to a standard protocol of an experimental research nursery (ONF-PNRGF, Pôle National des Ressources Génétiques Forestières, Cadarache, France). Seeds were collected from multiple *P. halepensis* trees within regional stands; consequently, the seedlings likely represent a mixture of genotypes (and potential maternal/epigenetic legacies), which reflects common nursery practice. Before planting, seeds were soaked in tap water for 24h and subsequently sown in the 1.4 L pots containing a substrate of 1:1 (v/v) RHP-certified blond peat and shredded bark of *Pinus pinaster* (5-10 mm in size), supplemented with mineral fertiliser 12-12-14 and 2 kg m^−3^ of Osmocote Exact 8/9 (ICL Growing Solutions, 15-9-11). Seedlings were initially grown in pots arranged in boxes holding 3 × 5 pots each for two months in a 300 m² greenhouse (March to April). Then, plants were grown for five months (May to September) outside of the greenhouse under optimal irrigation. Four weeks before the drought experiment, a selection of 120 seven-month-old saplings of similar size and vigor was moved to the greenhouse for acclimation.

### 2. Drought induction and experimental design

Before drought induction, soil field capacity (FC) was gravimetrically determined from the soil saturation point and gradual water loss curve. The 8-month-old plants were separated into 2 blocks, containing 4 boxes x 15 plants each (n = 60 per block). The first block consisted of control plants maintained at 60% FC, corresponding to hardening phase optimum. In the second block, drought was induced by stopping irrigation and reducing soil moisture until 40–45% field capacity (FC) was reached; this level was then maintained with reduced watering for approximately three weeks. At the end of the drought phase, plants were rewatered to 60% FC and monitored for four weeks to assess their recovery capacity (Figure 1).

**Figure 1.**
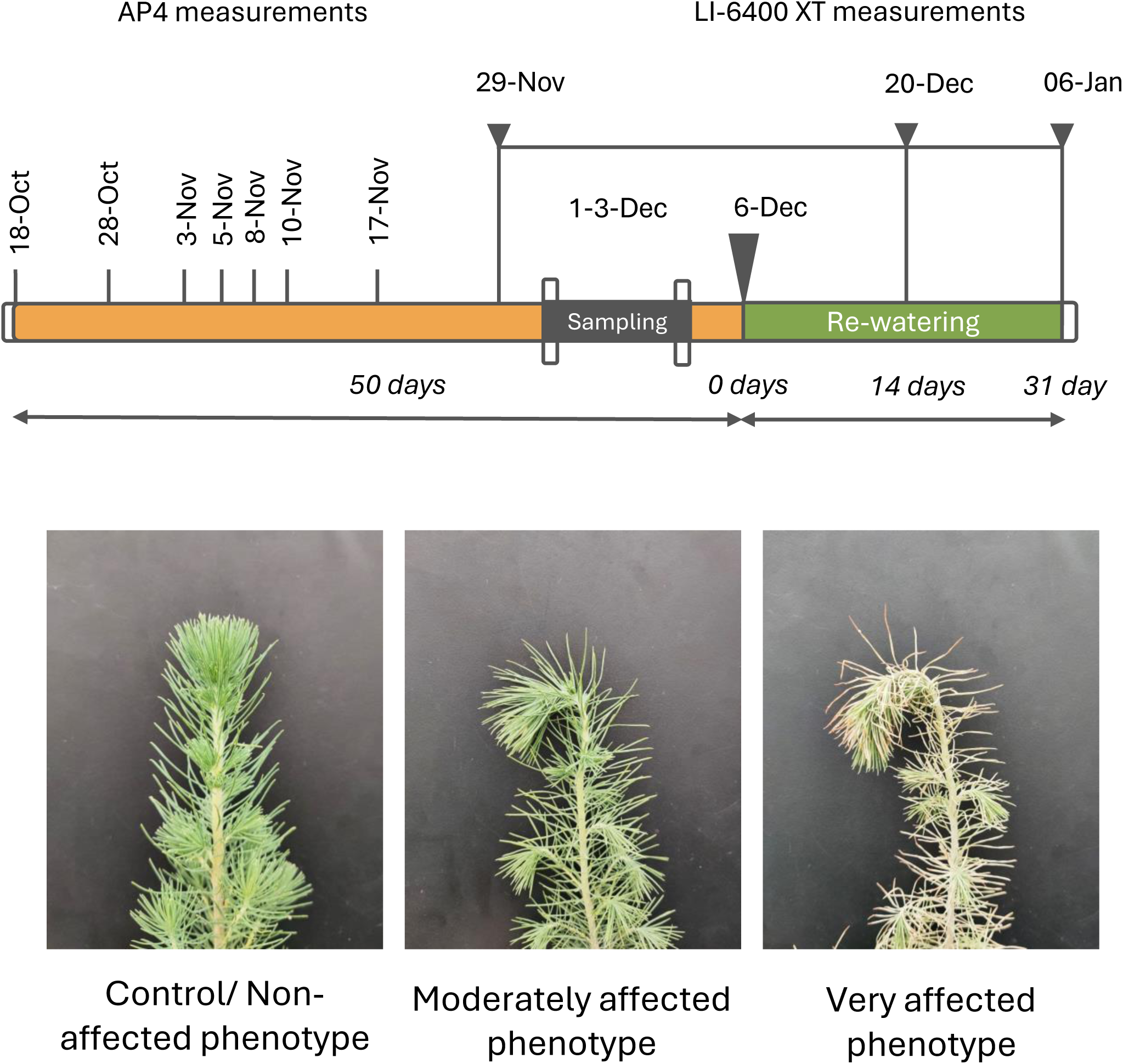
Experimental timeline, measurement schedule, and drought-induced phenotypes in the drought induction and recovery experiment with *Pinus halepensis*. The experiment consisted of three phases: drought treatment (orange, 50 days), sampling period (brown), and re-watering recovery (teal, 31 days beginning 6 Dec). Stomatal conductance measurements (AP4, black arrows) were taken at regular intervals during the drought phase (18 Oct - 17 Nov). Photosynthetic measurements (LI-6400 XT, purple triangles) were conducted at the end of drought period (29 Nov) and during recovery (20 Dec, 06 Jan). Plant and soil sampling (green triangle) occurred on 1-3 Dec during the sampling phase. Key physiological milestones included first drought symptoms at 31 days (18 Nov 2021) and 45% field capacity reached at 35 days (22 Nov 2021). Total experiment duration: 80 days (18 Oct - 06 Jan). Below are the pictures of drought-induced *P. halepensis* : non-affected control (CNA), non-affected drought (SNA), moderately affected with apical shoot wilting (SMA), and severely affected with apical shoot wilting, chlorosis, and needle loss (SVA).

### 3. Ecophysiological measurements

Stomatal conductance (gs) was monitored during the drying phase using a diffusion porometer (AP4, Delta T Devices, Ltd, Cambridge, UK). The instrument was calibrated to ambient humidity and temperature before each measurement round, with recalibration performed at ΔRh > 5% and Δ*T_amb_* > 2°C (Von Caemmerer and Farquhar, 1981). Measurements were made on single needles from the apex, alternating between control (n=8) and drought-treated plants (n=8).

For the recovery phase, combined net CO_2_ assimilation rate (*A_net_*) and stomatal conductance data from the apical shoot were obtained using a portable gas-exchange system with a transparent conifer chamber (LI-6400 XT, LI-COR Biosciences Inc., Nebraska, USA) under ambient photosynthetically active radiation (>300 µmol quanta m^-2^ s^-1^) and CO_2_ set to 400 ppm. The instrument was regulated to 20°C block temperature, flow to 500 ml min_-1_. Measurements took between 2-4 minutes, typically 3 minutes, to stabilize and were logged manually. Mean projected leaf area within the cuvette was determined as 39.9 cm² by scanning detached needles from apical shoots (n=4) and was used to scale measurements. Gas exchange parameters are calculated according to von Caemmerer and Farquhar 1981.

### 4. Plant and soil water content

Plants of different phenotypes (n = 8 per phenotype) were sampled at the end of drought induction phase (at 41 % FC), just before the onset of the recovery phase. Plant fine roots were cleaned from adhering soil by gently shaking, rinsing in MilliQ water, and blotting dry. Aliquots of 0.2 g were snap-frozen in liquid nitrogen, then -80°C until further analysis. Non-adhering root soil was collected and passed through 1.4 mm sieve to determine soil water content (SWC) according to the following formula:

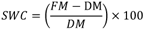

where *FM* is the fresh mass of the soil, *DM* is the dry mass (obtained by drying ∼20 ml of fresh soil at 70°C for 72 h). The same method was used to assess needle water content.

### 5. Nucleic acid extraction

Approximately 0.2 g of fresh roots were used for DNA and RNA extraction for metabarcoding studies. Because a single kit did not yield satisfactory results for both nucleic acids, the extraction protocol was optimized by using two commercial extraction kits: the ZymoBIOMICS DNA/RNA Miniprep Kit for RNA extraction and the FastDNA™ SPIN Kit for Soil (MP Biomedicals, USA) for soil DNA extraction. RNA purity was assessed by agarose gel electrophoresis and PCR amplification (*cf.* below) on 1 µL of RNA extract prior to reverse transcription. Then, RNA samples were reverse transcribed into cDNA using Transcriptor First Strand cDNA Synthesis Kit (Roche, Germany). Extracted nucleic acids was stored at -20°C. The quality and integrity of DNA and cDNA were evaluated using the Qubit dsDNA HS Assay Kit (Thermo-Fisher Scientific, Waltham, MA, USA). In addition, DNA and cDNA were tested for amplification using polymerase chain reaction (PCR). The V3–V4 variable regions of the 16S rRNA gene were targeted using the primers 515F (5’-TCG TCG GCA GCG TCA GAT GTG TAT AAG AGA CAG GTG YCA GCM GCC GCG GTA A-3’) and 806R (5’-GTC TCG TGG GCT CGG AGA TGT GTA TAA GAG ACA GGG ACT CAN VGG GTW TCT AAT-3’), each including Illumina adapter overhang sequences. Amplification was performed using the GoTaq® Flexi DNA Polymerase (Promega, USA, under the following conditions: initial denaturation at 95°C for 2 min; 34 cycles, each set at 94°C for 30 s, annealing at 53°C for 30 s, and extension at 72°C for 90 s, with a final elongation step at 72°C for 300 s.

### 6. Sequencing information

The V4 region of bacterial 16S rRNA genes were amplified using the primer set 515F Parada (5’-GTG YCA GCM GCC GCG GTA A-3’) and 806R Apprill (5’-GGA CTA CNV GGG TWT CTA A-3’). For fungal microbial profiling, the ITS2 region was amplified using the primers ITS3 (5’-GCA TCG ATG AAG AAC GCA GC-3’) and ITS4 (5’-TCCTCCGCTTATTGATATGC-3’). Amplification and purification of the libraries were carried out by Microsynth (Microsynth AG, Balgach Switzerland). Equimolar amounts of each library were pooled and submitted for sequencing. Sequencing was performed on the Illumina NovaSeq platform using the 250bp paired-end protocol. The resulting raw reads in FASTQ format were adaptor-trimmed, demultiplexed, and quality-checked.

### 7. Deep sequencing data processing

A total of 9,452,882 (fungal) and 9,771,280 (bacterial) demultiplexed reads were retrieved from the Illumina platform. Sequence data were analyzed using Qiime2 2023.5, (Bolyen et al., 2019), following quality filtering, primer removal, denoising, and chimera removal with the *dada2* plugin (Callahan et al., 2016). This process yielded 1,000,155 fungal and 1,087,779 bacterial high-quality sequences. Taxonomy was assigned to amplicon sequence variants (ASVs) using the *q2-feature-classifier* plugin (Bokulich et al., 2018) and the *classify-sklearn* naive Bayes taxonomy classifier, referencing the Greengenes2 16S rRNA database for bacteria (McDonald et al., 2024) and the UNITE v.9.0 rDNA ITS for fungi (Abarenkov et al., 2023).

Due to insufficient sequence depth (< 7302 for bacterial cDNA and < 2985 for fungal cDNA), one sample each dataset was excluded from further analysis. After removing mitochondrial and chloroplast DNA sequences, as well as singletons ASVs, 1,951 bacterial and 373 fungal ASVs were retained for downstream analysis. Subsequent analysis were performed in RStudio 2023.05.0 under R 4.4.2 (R Core Team, 2019). The Qiime2 outputs were imported to R using *qiime2R* package (Jordan E Bisanz, 2018) and then converted to *phyloseq* object format for further analysis (McMurdie and Holmes, 2013).

### 8. Microbiome data analysis

#### Alpha diversity

After rarefying to the minimal sample size, alpha diversity measures at the ASV level were calculated using the Shannon index (richness), inverse Simpson index (evenness) and phylogenetic diversity index (Faith’s PD). Statistical significance was assessed using the Welsh Anova test, followed by the Games-Howell post-hoc test (p < 0.05), both implemented in the *rstatix* R package (Kassambara, 2023).

#### Beta diversity

Community compositions were vizualised using PCoA ordination on TSS-normalized matrices based on weighted UniFrac distances (WUniFrac). Permutational analyses of variance (PERMANOVA) on the weighted UniFrac distance was performed to assess differences in microbiome composition matrices, using the *adonis2* in R (Oksanen et al. 2024). Additionally, pairwise PERMANOVA was conducted to identify differences in bacterial community composition between the different treatment-phenotypes (Arbizu, 2020). The significance level was set at p < 0.05 for PERMANOVA and False Discovery Rate (FDR) < 0.05 for pairwise PERMANOVA analysis.

#### Phenotype-microbiome association test

A Distance-Based Microbiome Kernel Association Test With Multi-Categorical Outcomes (MiRKAT-MC) was employed to assess the association between microbiome community composition and treatment-phenotype variables based on omnibus of dissimilarity matrices (“Jaccard”, “BC”, “U.UniFrac”, “G.UniFrac.00”, “G.UniFrac.25”, “G.UniFrac.50”, “G.UniFrac.75”, “W.UniFrac”). To ensure the robustness of associations across different mathematical assumptions about sample relationships, analyses were performed using four kernel types: Linear Kernel (LK) for proportional relationships, Gaussian Kernel (GK) for smooth neighborhood patterns, Exponential Kernel (EK) for moderate decay functions, and Laplacian Kernel (LaK) sensitive to sharp similarity boundaries. Phenotypes were classified in three groups, “non-affected’, ‘moderately affected’ and “severely affected’ classes, ordered by severity. MiRKAT-MC was chosen due to its ability to account for complex interactions and associations within microbiome data, making it suitable for exploring links between microbial community structure and treatment-phenotype effects. (Jiang et al., 2022a; Sun et al., 2023)

#### Differential abundance analysis

Given the inherent compositional structure of microbiome count data, Analysis of Compositions of Microbiomes with Bias Correction 2 (ANCOM-BC2) was employed due to its capacity to correct for compositional effects and sample-specific biases, ensuring more accurate identification of differentially abundant taxa (Lin and Peddada, 2020).

#### Ecological role and function prediction

Prior ecological functions prediction of taxa, bacterial and fungal ASVs were prevalence filtered (≥ 5% prevalence, detection threshold of 4). Bacterial functions were predicted using FAPROTAX v.1.2.7 (Louca, 2022). Functional prediction of fungal guilds and functions was performed using FUNGuild and FungalTraits databases, with assignments carried out. in *microeco* R package (Liu et al., 2021). Both tools rely on trait-based inference from taxonomic assignments and are limited by the completeness of their reference databases. Functional predictions should therefore be interpreted as indicative rather than definitive, and do not replace direct measurement of functional activity. Overall, bacterial functions were predicted for 48.0% of ASVs, while fungal functions were assigned to 75.5% with FUNGuild and 64% with FungalTraits. Read counts were normalized to the total number of functional matches and to the total read abundance per sample. Statistical testing was done using the non-parametric Welch’s ANOVA test, setting the significance level at p < 0.05, followed by a post-hoc Dunn’s test with FDR correction for multiple comparisons using the *rstatix* package (Kassambara, 2023).

#### Inferring Community Assembly Mechanism

Investigation of the microbial and fungal community assembly processes and their relative importance was estimated using phylogenetic bin-based null modeling approach, implemented in the iCAMP v1.5.12 R package (Ning et al., 2020). Assembly processes were quantified using null model analysis based on beta mean nearest taxon distance (bMNTD) and taxonomic β-diversities (Ning et al., 2020) using the Confidence significance test to evaluate how the observed beta diversity indices deviate from null expectation for each phylogenetic bin. The bin parameters —minimum bin size and maximum phylogenetic distance—were set at 24 and 0.2 for bacteria and 10 and 0.2 for fungi, following recommendations from the original study. This framework partitions ecological processes in the following selection groups: HoS, homogeneous selection; HD, homogeneous dispersal; DR, ecological drift; DL, dispersal limitation; HeS, heterogeneous selection.

#### Interaction networks

Microbial Association Networks were constructed to uncover inter- and intra-phylum interaction patterns among treatment phenotypes. Prior to network construction, bacterial and fungal ASVs were filtered based on prevalence (≥ 5%) and a detection threshold of 4. Microbial co-occurrence networks were constructed using SparCC algorithms from the SpiecEasi package to correlate bacterial and fungal species interactions at the genus level, and retaining only significant associations (|r| ≥ 0.7, FDR < 0.05) using *microeco* (v1.8.0) (Liu et al., 2021). Network modules were identified using fast greedy algorithm (Clauset et al., 2004). For network characterization and comparison, network parameters and topological properties were calculated using the *meconetcomp* package (v 0.5.1) (Liu et al., 2023). Within-module connectivity (Zi) and among-module connectivity (Pi) metrics were used to classify node roles. Network data was exported and visualized using Gephi software v.0.10.0 (Bastian et al., 2009).

#### Data Availability Statements

Data and code supporting the findings of this study will be made publicly available upon acceptance. Sequencing data and associated metadata will be deposited in Zenodo, and analysis scripts will be available on GitHub.

#### Generative AI Usage Statement

During the preparation of this work, the authors utilized Claude Sonnet 4.6 solely for grammatical correction and language refinement. After using this tool/service, the authors carefully reviewed and revised the content where necessary and take full responsibility for the final version of the publication.

## III. Results

### 1. Impact of drought and subsequent re-watering on foliar gas exchange and visual symptoms

In mid-October, watering was stopped to induce progressive soil drying and hardening of the plantlets. After 15 days, soil moisture in the control treatment reached 60% of field capacity (FC). From that point onward, maximum and minimum temperatures remained between approximately 5°C and 25°C, enabling the establishment of drought stress conditions without the confounding influence of high temperatures (Figure 2A). In the drought treatment, soil moisture continued to decline and stabilized at 45% FC after 35 days (Figure 2B). Stomatal conductance (*g_s_*) in drought-treated plants began to diverge from control plants at 56% FC (9% lower than in controls), with *g_s_* decreasing by almost 30% from 370 to 270 mmol m⁻² s⁻¹ (Figure 2C). Once the target FC of 45% in the drought treatment was attained, *g_s_* was three to four times lower in the drought as compared to the control group. Drought treatment significantly affected plant growth, leading to a reduction in shoot diameter (-5.77%) and growth rate (2.4% in control vs. 0.7% in drought treatment) (Supplementary Figure 1; Yuen trimmed means test, p < 0.05).

**Figure 2.**
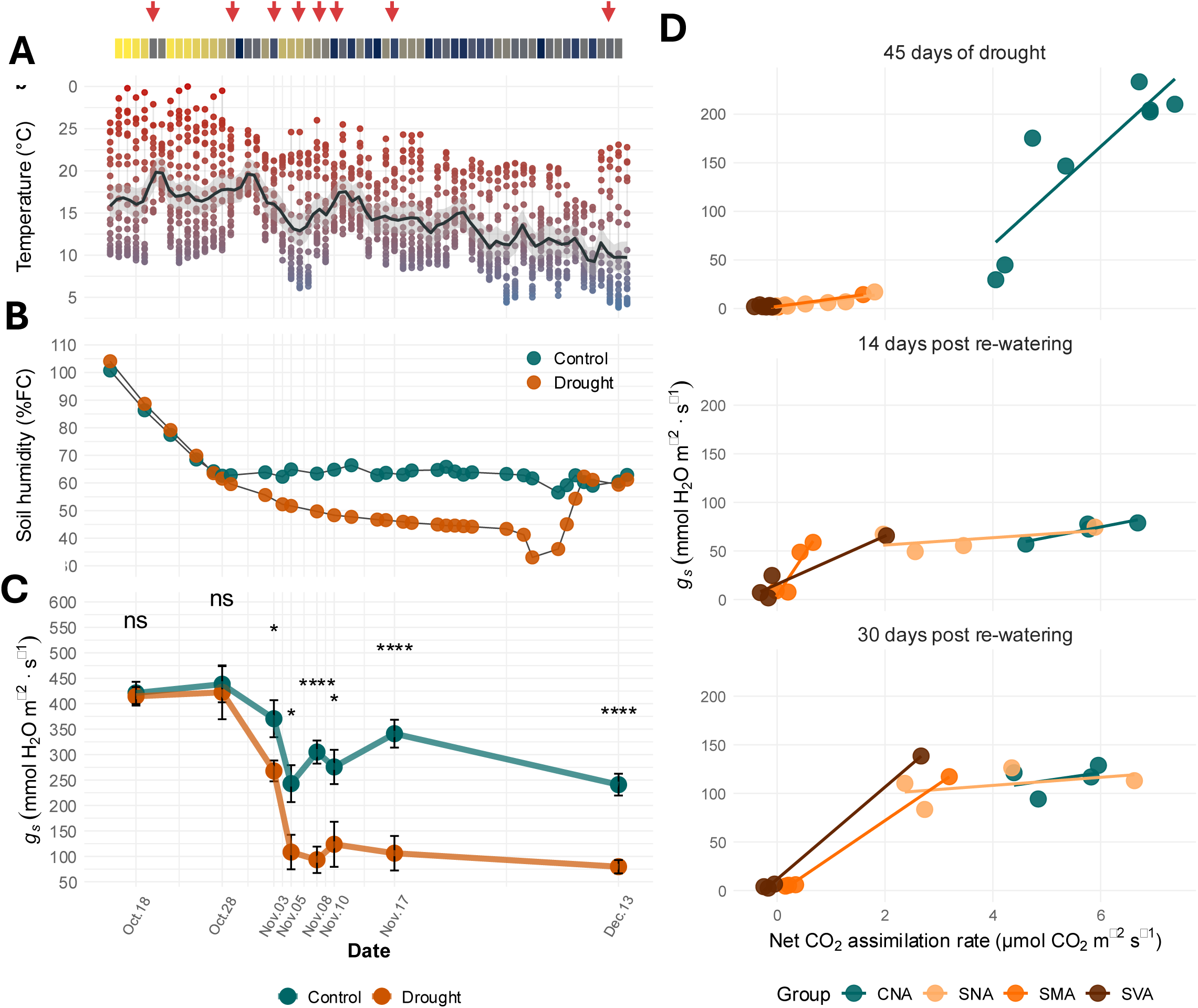
Environmental conditions, physiological responses, and recovery patterns during drought stress experiment. **(A) G**reenhouse luminance (upper panel) throughout the experiment and daily temperature (lower panel) variation throughout the experiment with measurement dates indicated by red arrows. orange arrows indicate the specific days on which measurements shown in Figures 2b and 2c were conducted. **(B)** Soil humidity (%FC) in control and drought treatments. **(C)** Stomatal conductance (*g*s) responses at specific measurement dates in control and drought treatments. **(D)** Relationship between net CO_2_ assimilation rate and stomatal conductance at three time points: 45 days of drought (top), 14 days post re-watering (middle), and 30 days post re-watering (bottom). Plants were categorized by visual symptoms in control (C) and drought-submitted (S) groups as follows: non-affected (CNA and SNA; for control and S groups, respectively), moderately affected with apical shoot wilting (SMA), and severely affected with apical shoot wilting, chlorosis, and needle loss (SVA). Lines represent linear regression fits for each symptom category. Error bars represent mean ± SE. Statistical significance determined by Welch’s t-test (ns = not significant, * *p* < 0.05, **** *p* < 0.0001). Sample size *n* = 8 per treatment for panel **(C)** and n = 4-8 for panel **(D)**.

Along drought treatment and the rewatering, we assessed visual phenotypes across treatments (Figure 1). The observed symptoms were categorized as four treatment-phenotypes: ‘non-affected’ (CNA and SNA, for the control (C) and drought treatment, respectively, with “S” indicating ‘submitted to drought’), ‘moderately affected’ with apical shoot wilting (SMA), and ‘se<u>v</u>erely affected’ with apical shoot wilting, chlorosis, and needle loss (SVA). Forty-five days after beginning the drought experiment, as well as 2 weeks and one month after rewatering, we conducted combined CO_2_ and H_2_O gas exchange analyses on the apical shoot as to determine whether these were linked to the visual phenotypes (Figure 2D).

Stomatal conductance in moderately and severely affected plants (SMA, SVA) was very low, with net CO_2_ assimilation rate (*A_net_*) mostly negative, indicating that respiration dominated. This was less pronounced in the drought-treated plants without symptoms (SNA), which showed positive *A_net_* values at relatively low stomatal conductance. Nonetheless, their physiological responses were more comparable to those of moderately (SMA) and severely (SVA) affected plants, despite their phenotype resembling that of the control group (CNA) (Figure 2D, upper panel). In the CNA group, a wide range of *g_s_* and *A_net_* values was found. Although unexpected, given that irrigation was strictly controlled across treatments (confirmed by box mass measurements of 15 pots; (Supplementary Figure 2), variations in soil water content were observed among individual pots at harvest. This localized variability in soil water availability directly corresponded to the emergence of distinct drought phenotypes, as evidenced by their specific ranges in stomatal conductance (*g_s_*).

Rewatering had strikingly binary effects on gas exchange recovery, which became even more pronounced 30 days after rewatering than after 14 days (Figure 2D, middle and lower panels). A clear uncoupling between stomatal conductance (*g_s_*) and net CO_2_ assimilation (*Aₙₑₜ*) was observed. Nearly half of the plants (mostly SNA) rapidly regained *g_s_* levels comparable to the control group, although only about 10% fully recovered *Aₙₑₜ*. Only about 1/4 recovered to 50% of the control *Aₙₑₜ* levels. The remaining symptomatic plants maintained low *g_s_* values with more than one order of magnitude lower than the control and recovered plants. Moderately affected plants (SMA) showed slightly higher *Aₙₑₜ* than severely affected ones (SVA), which continued to exhibit negative values (Figure 2D, middle and lower panels). Below, the resident and active soil microbiome of these plants are investigated for the end of the drought period.

### 2. Association of drought-induced *P. halepensis* phenotypes with specific microbial and fungal diversity patterns

Drought affected both the resident (gDNA) and active (cDNA) bacterial communities, in a somewhat unexpected way. In drought-treated plants, alpha diversity of the resident bacterial microbiome increased compared to controls, in terms of both richness and evenness (Shannon and Inverse Simpson indices, respectively; p < 0.01), as well as phylogenetic diversity (Faith’s PD, p < 0.01). Notably, these differences were more pronounced in the resident fraction, than in the active fraction, where richness was unaffected by drought, but evenness and phylogenetical diversity were significantly altered. Interestingly, the most pronounced differences in both fractions were between CNA, SMA and SVA, respectively (Games-Howell, p < 0.05). Remarkably, the alpha diversity of SNA was comparable to that of SMA and SVA but significantly differed from control plants of the same phenotype (CNA), in terms of richness and evenness within the resident fraction (Games-Howell, p < 0.05) (Figure 3A).

**Figure 3.**
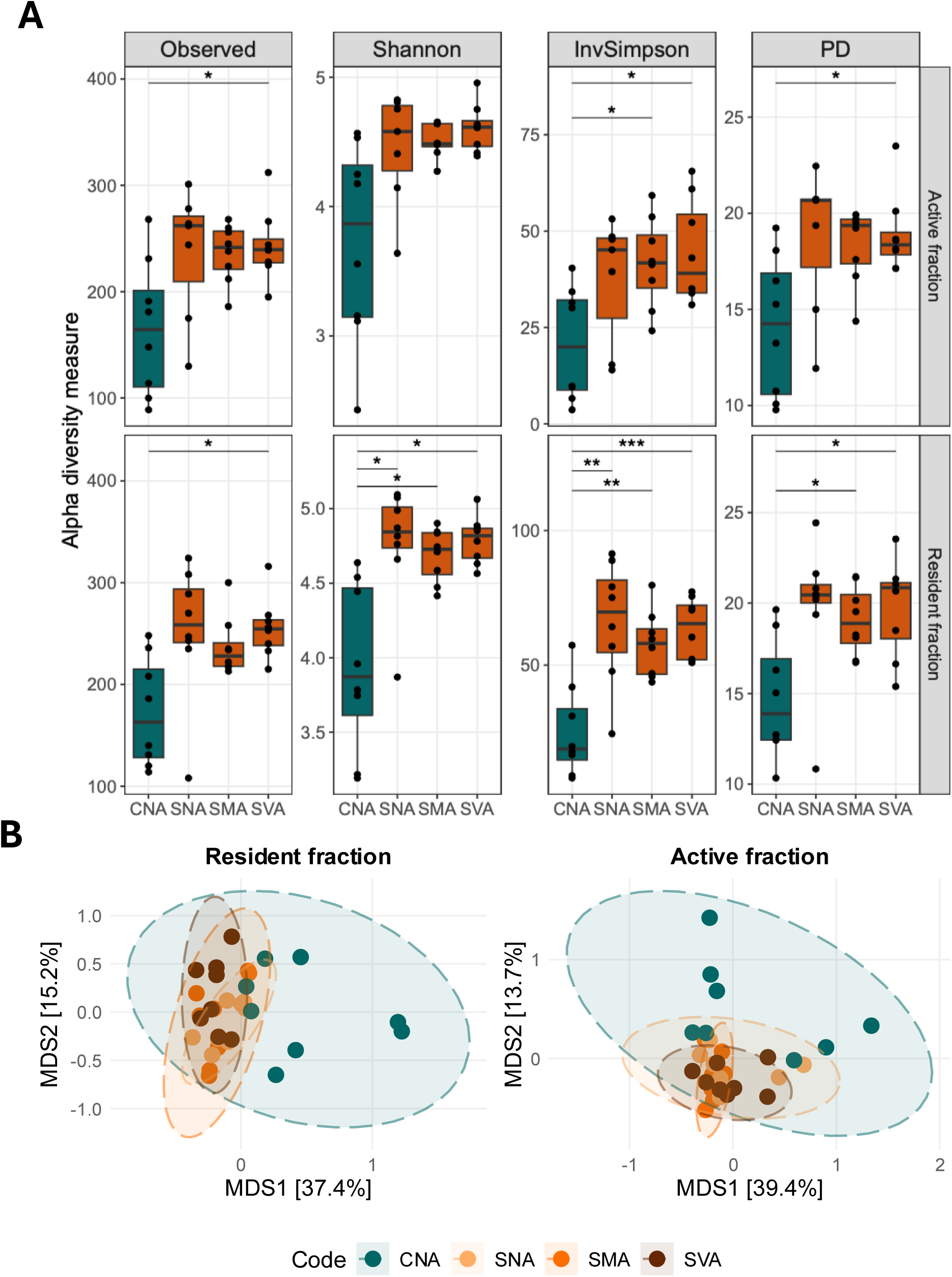
Microbial diversity associated with *Pinus halepensis*. **(A)** Alpha diversity metrics showing significant differences between treatments (Welch ANOVA test). Pairwise comparisons were conducted using Games-Howell test with significant results indicated. **(B)** Beta diversity analysis using multidimensional scaling (MDS) of weighted UniFrac distances for active and resident microbial fractions. Statistical significance: * *p* < 0.05, ** *p* < 0.01, *** *p* < 0.001, **** *p* < 0.0001.

In contrast, fungal alpha diversity metrics were not affected by drought, showing similar levels of richness, evenness, and phylogenetic diversity to well-watered control plants (Supplementary figure 3A).

Microbiome composition ordination using PCoA on weighted Unifrac (WUniFrac) distances explained a substantial fraction of the variance, for both microbial and fungal communities - 52.6% and 56.9% in the resident fraction and 53.1% and 55.6% in the active fraction across the two first principal axes (Figure 3B and Supplementary figure 4).

We then investigated whether the community structure of root-associated bacterial and fungal communities differed among treatment-phenotypes, considering both resident and active fractions. The bacterial community structure was significantly influenced by the water regime in both fractions (p < 0.001), but not by phenotype alone (p > 0.05) (Table 1). However, the interaction between water regime and phenotype explained the largest portion of the variance: 29.8% in the resident (gDNA) fraction and 22.0% in the active (cDNA) fraction (Table 1). Additionally, the CNA group significantly differed from SNA, SMA and SVA in the resident fraction (pairwise PERMANOVA, p.adj < 0.01), a pattern that was also observed in the active fraction (pairwise PERMANOVA, p.adj < 0.05).

**Table 1.**
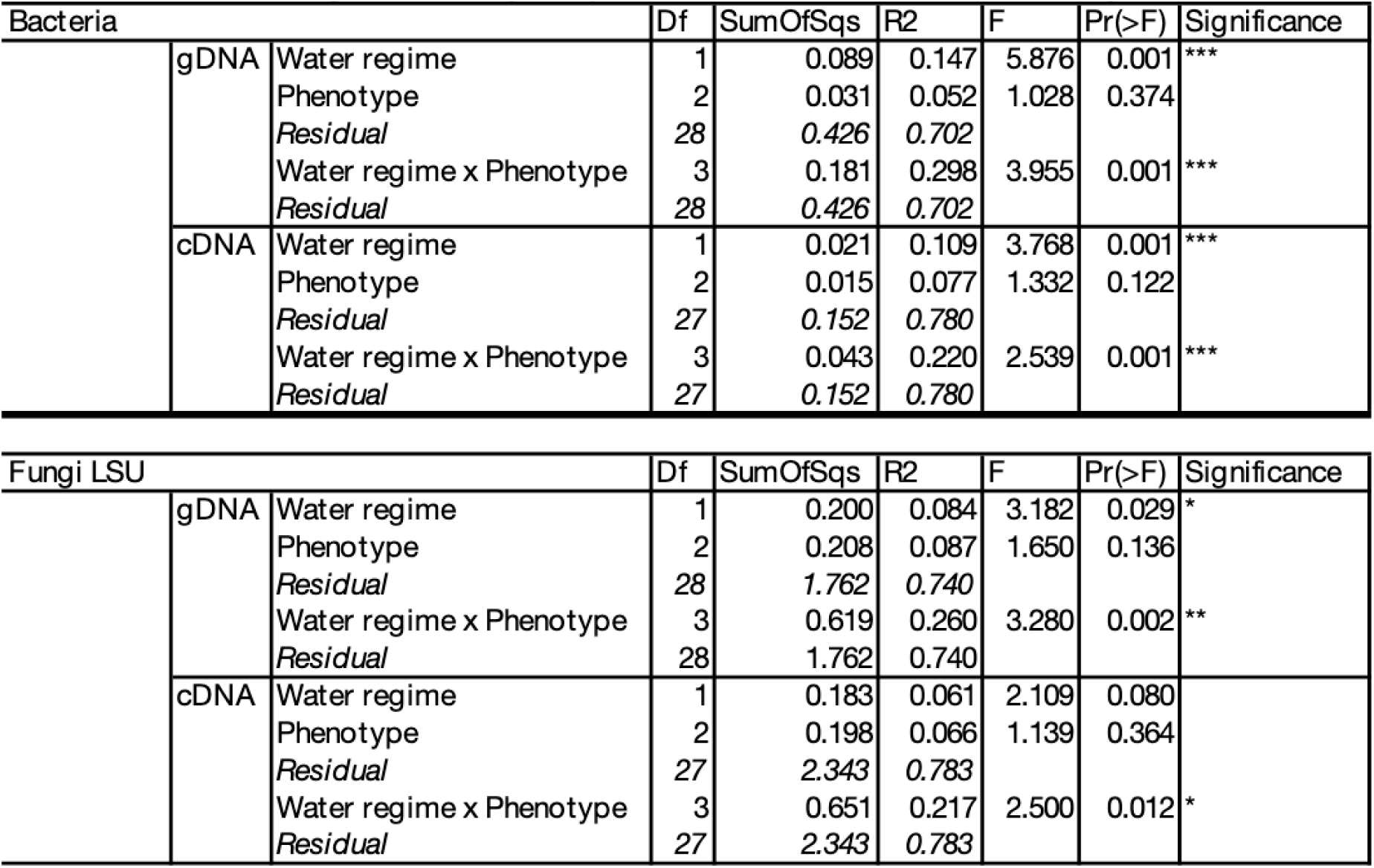
PERMANOVA results showing the effects of water regime and phenotype factors on bacterial and fungal community composition based on Wunifrac dissimilarity.

In the fungal community, a similar pattern was observed for the resident fraction: although the water regime had a significant effect (p < 0.001), the largest portion of the variance was explained by its interaction with phenotype (26%, p < 0.002) (Table 1). In the active fraction, the interaction between water regime and phenotype was the only factor that significantly explained the variance (21.7%, p < 0.012) (Table 1).

### 3. Drought-induced specific phenotype association strength with root microbiome

To test the strength of association between drought-induced phenotypes and microbiome composition, we applied the Distance-Based Microbiome Kernel Association Test for Multi-Categorical outcomes (MiRKAT-MC). This method uses a semi-parametric kernel machine regression model to test for associations between microbiome composition (represented by a kernel matrix) and multivariate categorical outcomes (ordinal or nominal (Jiang et al., 2022b).

The strongest microbiome-phenotype association was detected in cDNA, with generalised phylogenetic distance metrics (Generalised UniFrac with 0 < α < 0.75) showing the highest confidence (p < 0.001). Non phylogenetic distances yielded comparable results, where Jaccard index showed highly significant results across all kernel types (p < 0.001), as did the Bray-Curtis dissimilarity (p < 0.01). Interestingly, WUniFrac showed less significant result compared to other metrics (p < 0.05).

In the gDNA dataset, all distance metrics tested showed significant associations. Both WUniFrac (p = 0.002-0.004 across kernels) and unweighted UniFrac (p = 0.020-0.029) showed significant associations with phenotype. Generalized UniFrac (G.UniFrac) metrics were also significant, with p-values decreasing as the weight increased (G.UniFrac.00, p = 0.017-0.022; G.UniFrac.75, p = 0.002-0.004).

Overall, the MiRKAT-MC omnibus test revealed significant associations for fractions, with particularly strong results for the active (p = 2.9 × 10⁻⁵) compared to the resident fraction (p = 6.8 × 10⁻³).

### 4. Inferring microbiome community assembly mechanism associated with drought induced phenotypes

After demonstrating that phenotype–microbiome associations were primarily mediated by abundance–phylogeny relationships, we sought to explore how these associations related to assembly mechanisms of microbial communities. To this end, we employed iCAMP (Ning et al., 2020), an approach that integrates taxonomic, phylogenetic, and ecological data to quantify the relative contributions of deterministic and stochastic processes.

In the resident fraction, most taxon turnover was attributed to stochastic assembly process (> 60% across all treatment-phenotypes), with drift accounting for the largest part (39-47%). Deterministic process turnover showed a phenotype-dependent pattern, with CNA and SNA (control and drought-submitted non-affected plants) showed 29% and 30.5% deterministic assembly, respectively while moderately and severely affected plants (SMA and SVA) showed increased contributions of deterministic processes (34.9% and 37.2%, respectively), indicating a link between symptom severity and deterministic microbiome assembly.

In the active fraction, deterministic taxon turnover was more strongly associated with water regime than with phenotype, with CNA plants exhibiting 19.6% deterministic turnover compared to 35.5%, 32.2%, and 30.8% for SNA, SMA, and SVA plants, respectively. Remarkably, drift was least important in SNA plants (31.1%), while dispersal limitation (DL) showed similar contributions between CNA and SNA plants (28.7% and 27.6%, respectively) (Figure 4A).

**Figure 4.**
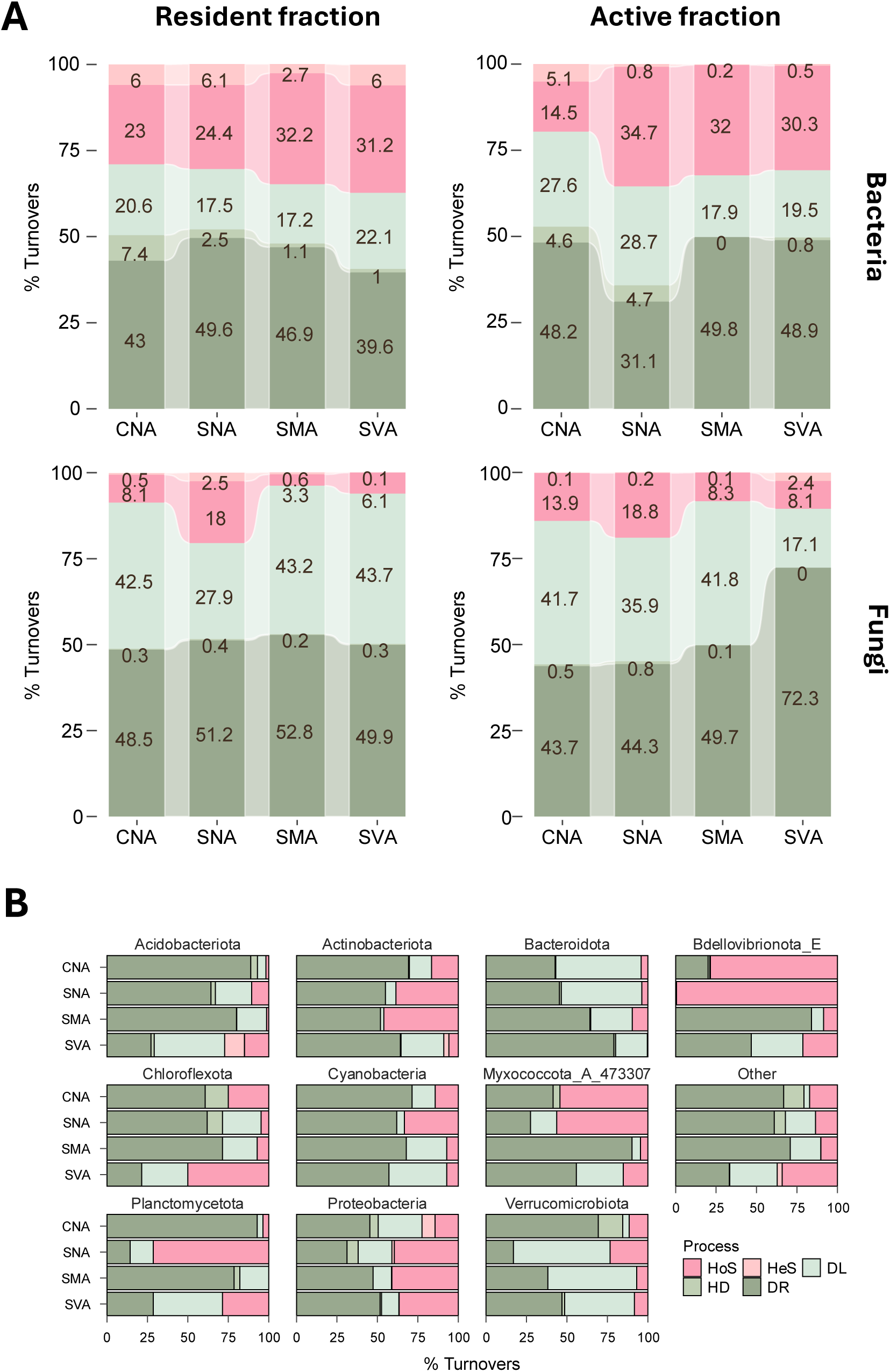
Ecological assembly mechanisms of resident (gDNA-based) and active (cDNA-based) microbiomes associated with *Pinus halepensis* treatment-phenotypes. **(A)** Relative contributions of community assembly processes within each treatment-phenotype **(B)** Assembly mechanism of the top ten the most abundant bacterial taxa. Analysis was conducted using the iCAMP R package, to infer the dominant ecological drivers across different treatment-phenotypes. **DL**, dispersal limitation; **DR**, drift; **HD**, homogenizing dispersal; **HeS**, heterogeneous selection; **HoS**, homogeneous selection.

We further investigated the assembly processes of the most abundant phyla in both gDNA and cDNA datasets. In SNA plants, deterministic turnover under homogeneous selection was attributed to *Bdellovibrionota_E*, *Planctomycetota* and *Myxococcota_A_473307*, collectively accounting for more than 50%. Interestingly, the deterministic turnover of *Actinobacteriota* peaked in SNA and SVA plants (approximately 45-50%) and was the lowest in SVA within the active fraction (Figure 4B). In gDNA *Bdellovibrionota_E* showed less deterministic turnover in CNA and SNA (Supplementary figure 5).

Interestingly, fungal community turnover followed a different pattern. In the resident fraction, deterministic turnover spiked dominated only in SNA (20.5 %) (Figure 4A). In the active fraction, the contribution of drift (DR) increased sharply from 43.7.6% in CNA to 72.3. % in SVA, while dispersal limitation (DL) declined in parallel from 41.7% to 17.1% (Figure 4A).

Overall, bacteria and fungi showed contrasting assembly dynamics. In the bacterial community, deterministic processes became more dominant with increasing drought severity and symptom progression. Conversely, fungal communities shifted toward stochastic assembly under drought, with a marked rise in drift, particularly in the severely affected phenotypes.

### 5. Co-occurrence network analysis

Co-occurrence network analysis was used to explore topological properties and taxonomic (at the level of genus) interactions in bacterial and fungal communities, and how these were impacted by drought across the phenotype gradient.

The CNA network was the most complex, containing more nodes (vertex) and edges than the SMA, SNA, and SVA networks. Despite its larger size, the CNA network exhibited lower density than the others, indicating a more loosely connected structure. It also showed the highest heterogeneity, suggesting greater variability in node connectivity, while maintaining similar centralization compared to other networks. Drought-induced phenotypes, by contrast, exhibited sparser and less compact network structures than well-watered plants, consistent with a drought-induced disruption of microbial network connectivity. Notably, SNA network showed stronger local organization and community structure, despite having fewer overall connections (Figure 5A, Table 2).

**Figure 5.**
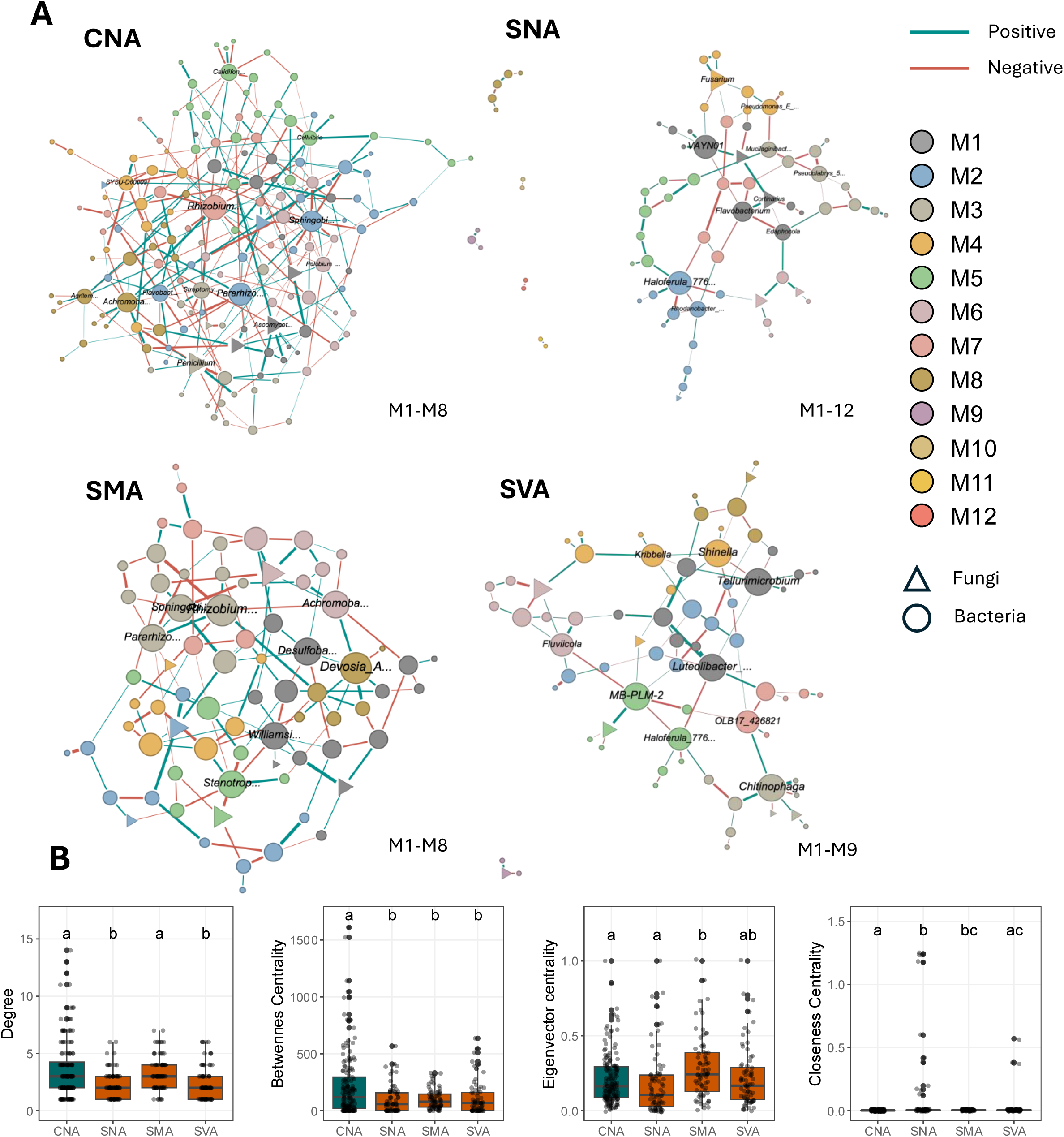
Co-occurrence networks between bacterial and fungal genera. **(A)** Networks of SparCC correlations between fungal and bacterial species. Nodes represent microbial taxa coloured by network module, with fungi shown as triangles and bacteria as circles. Edges indicate significant correlations between taxa. Node size represents degree centrality. Edge colour indicates correlation sign (positive/negative), and edge width represents correlation strength. **(B)** Calculated network parameters and topological properties. Letters represent pairwise comparison results from Welch ANOVA with Games-Howell post-hoc test (p.adj < 0.05).

**Table 2.** Network topological parameters.

| Parameters | CNA | SNA | SMA | SVA |
| --- | --- | --- | --- | --- |
| Vertex | 156 | 74 | 74 | 72 |
| Edge | 281 | 84 | 124 | 90 |
| Average degree | 3.6 | 2.27 | 3.35 | 2.5 |
| Average path length | 2.96 | 3.98 | 3.04 | 3.54 |
| Network diameter | 6 | 9 | 7 | 8 |
| Clustering coefficient | 0.02 | 0.06 | 0.04 | 0.01 |
| Density | 0.02 | 0.03 | 0.05 | 0.04 |
| Heterogeneity | 0.72 | 0.54 | 0.44 | 0.61 |
| Centralization | 0.07 | 0.05 | 0.05 | 0.05 |
| Modularity | 0.53 | 0.7 | 0.54 | 0.64 |

Degree distributions were similar between CNA and SMA, whereas SNA and SVA networks exhibited lower node interconnectivity. Betweenness centrality was higher in CNA compared to lower and similar values in SNA, SMA, and SVA. Closeness centrality displayed the greatest variability in SNA, and remained relatively low and uniform under other conditions. (Figure 5B). Interestingly, eigenvector centrality was the lowest in CNA, indicating a more distributed influence pattern, whereas it was higher in SNA and SMA, suggesting a more hierarchical structure with influence concentrated among a subset of highly connected taxa (Figure 5B).

To assess individual taxa roles in network topology, we examined node hubs and connectors. The CNA network included three node hubs, *Calidifontimicrobium, Flavobacterium*, and unidentified *SYSU-D60009*, which showed strong within-module importance. Remarkably, *Rhizobium* played a keystone role as a network hub in CNA network structure, but was nearly absent in drought-induced phenotypes, suggesting its importance in maintaining microbial network structure under well-watered conditions. In SNA, SMA, and SVA networks, nodes tended to act as peripheral elements with localized connections or as connector nodes linking different modules. Notably, only in SMA did a fungal taxon (unidentified Ascomycota) served as a connector; in all other drought phenotypes, fungi remained peripheral, indicating a limited role in stabilizing microbial networks (Figure 5; Supplementary figure 6).

We further compared node sources of edges across networks. Positive associations were most frequent within Proteobacteria and between Proteobacteria and Bacteroidota. Conversely, negative associations were primarily observed within Proteobacteria, between Proteobacteria and Bacteroidota, and between Proteobacteria and either Verrucomicrobiota or Actinobacteria. Although negative interactions involving Ascomycota and bacterial phyla (Proteobacteria, Bacteroidota) were present, they represented only a minor fraction (Supplementary figure 7).

Analysis of node and edge conservation across treatment-phenotypes revealed striking divergence. The CNA network had the highest number of unique nodes, sharing only 3.5% with SNA, 8.8% with SMA, and 3.5% with SVA. Remarkably, almost no edges were shared between treatment-phenotypes, suggesting a near-complete rewiring of microbial interactions under drought, despite some shared taxa (Supplementary Figure 7).

### 6. Taxa ecological and functional characterization

According to FAPROTAX annotations, the most abundant bacterial functional groups across both active and resident fractions were aerobic and anaerobic chemoheterotrophs. A substantial number of taxa were assigned to nitrogen cycling, particularly ureolysis (2.1-3.8%), nitrogen fixation (1.7-3.7%), nitrate respiration (0.7-3.6%), and nitrate reduction (0.9-4.7%) (Figure 6A).

**Figure 6.**
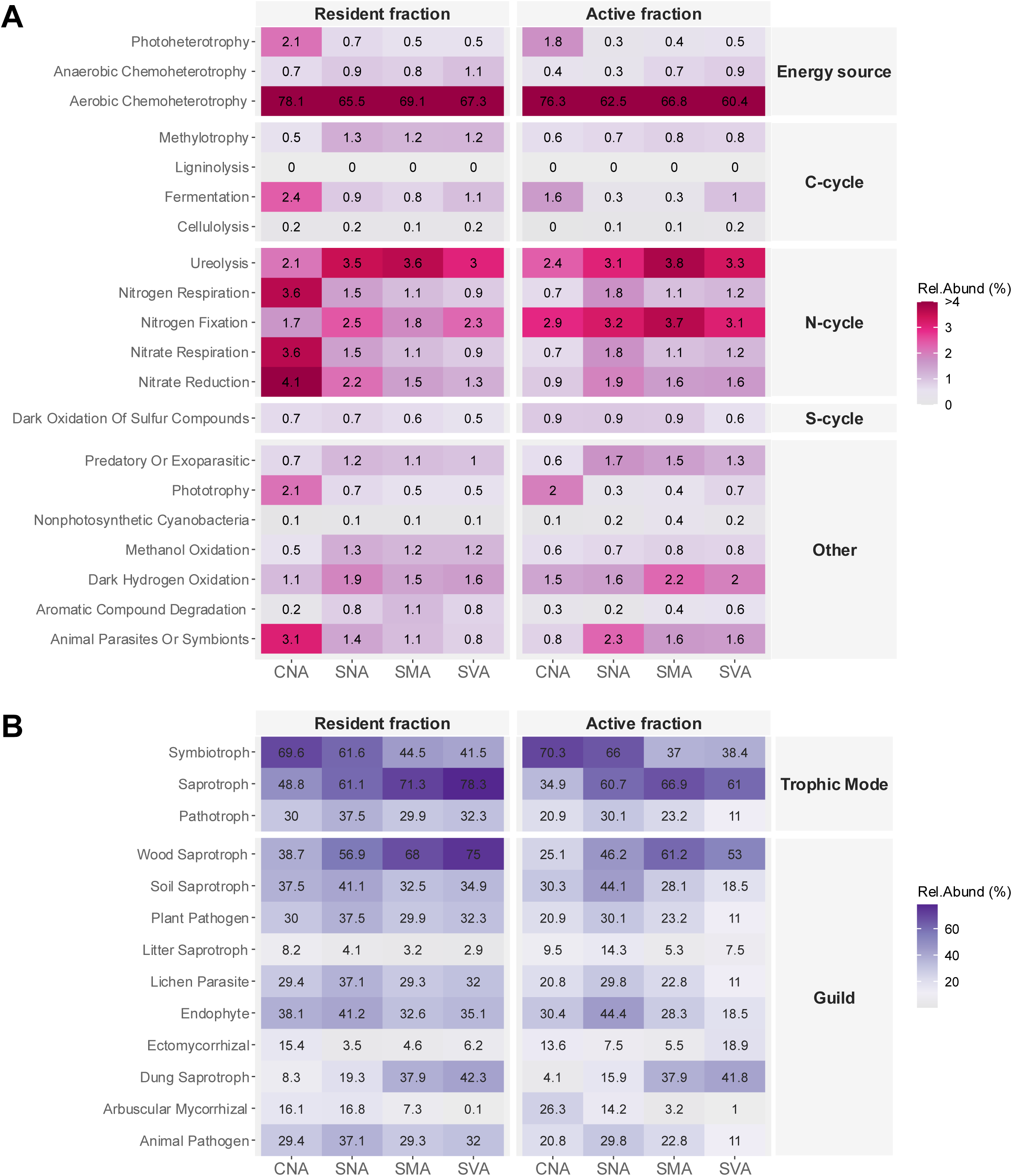
Functional annotation of microbial and fungal communities based on taxonomical parsing. A. Taxonomic parsing of bacterial communities based on FAPROTAX functional database. B. Taxonomic parsing of fungal taxa based on FUNGuild functional database. Heat map displays the relative abundance of predicted metabolic functions in active (cDNA) and total (gDNA) fractions across control (CNA) and drought-induced phenotypes (SNA, SMA, SVA). Values represent the percentage of ASVs assigned to each functional category, weighted by the relative abundance of each taxon.

In both active and resident fractions, the proportion of ureolysis-capable bacteria was higher in drought-induced phenotypes (SNA, SMA, and SVA) compared to the control group (CNA). The abundance of nitrogen-fixing ASVs was also elevated under drought, with the most pronounced increase observed in the active fraction (cDNA). Interestingly, the highest proportions of nitrate reduction and nitrate respiration were observed in CNA within the total community (gDNA) (Figure 6A).

For fungi, ecological function was assigned using FUNGuild to classify taxa by trophic mode and guild. Most taxa were assigned to symbiotrophic lifestyle, representing 41-73% in the active fraction and 41-68% in the resident fraction. The dominant fungal guilds were wood saprotrophs (37-76% in the resident and 20-52% in the active fraction) and soil saprotroph (active 21-38%, resident 32-42%).

Interestingly, the proportion of saprotrophic fungi increased under drought SNA, SMA SVA) indicating that drought promoted growth and/or physiological activities of these fungi. In addition, the proportion of symbiotrophs declined in drought-induced phenotype (Figure 6B).

### 7. Differential abundance analysis

To identify drought-resistant taxa associated with the non-affected phenotype, we conducted a differential abundance analysis between plants of the same phenotype under different water regimes (“non-affected”, CNA and SNA) (Figure 7). Several bacterial genera, including Calidifontimicrobium, Arachidicoccus, an unresolved taxon (*Rhodoferax_A_585629*), and *Pseudomonas_E_647464* (order level) were significantly more abundant in SNA compared to CNA (p < 0.05). Similarly, several fungal genera were enriched in SNA, particularly *Lilapila, Mortierella, Trichoderma, Cortinarius*, and *Rhizophydiales_gen_Incertae_sedis* (order level) (p < 0.05).

**Figure 7.**
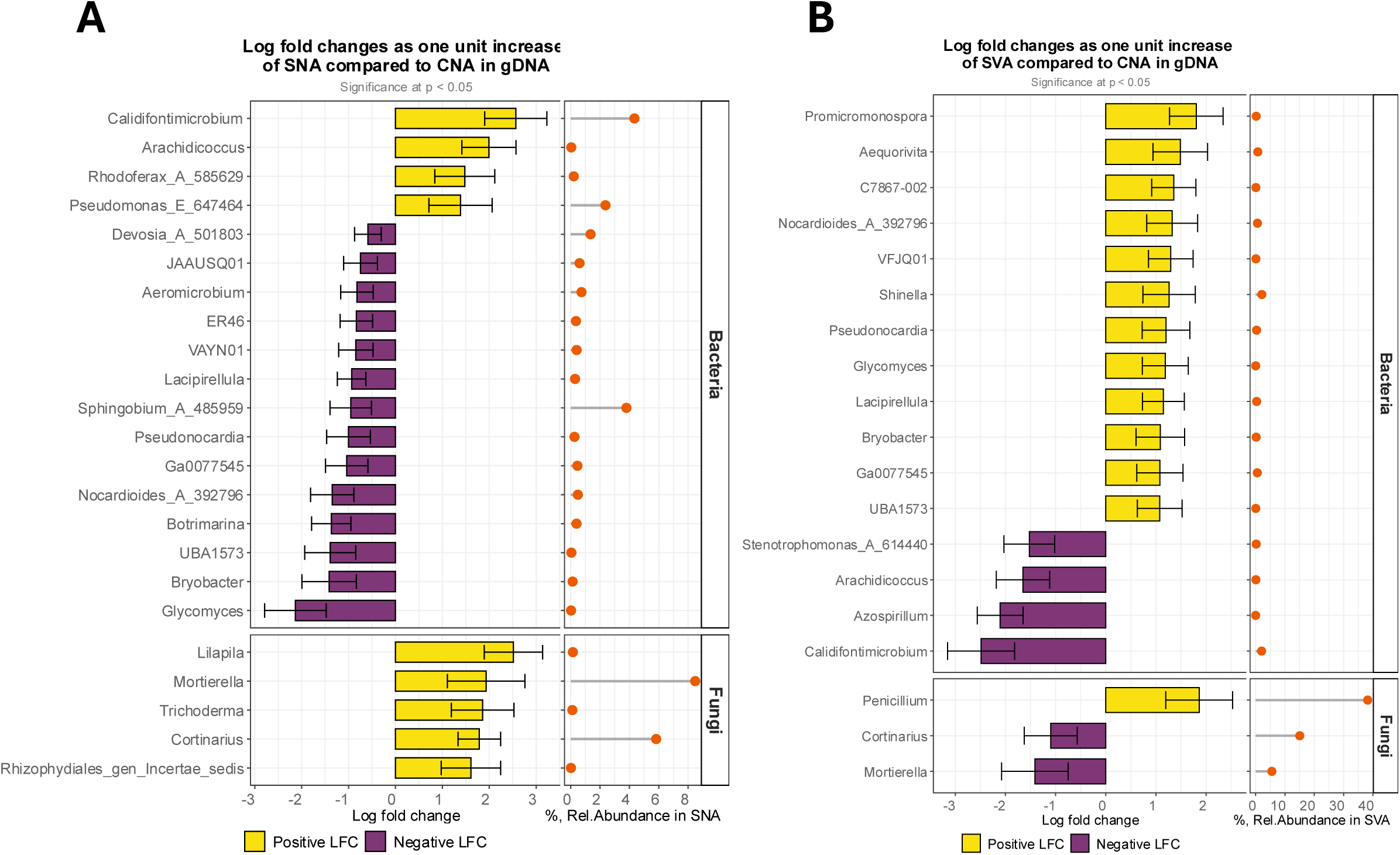
Differential abundance analysis *Pinus halepensis* roots at Genus level between selected treatment phenomes. (A) SNA and CNA (B) SVA and CAN. Analysis was performed using ANCOM-BC2, significant result showed at p < 0.05.

Despite sharing the same phenotype, some genera were negatively affected by drought, with reduced abundance in SNA. This included, among other genera, *Nocardioides_A_392796, Bryobacter, UBA1573, Glycomyces, and Pseudonocardia*.

When comparing two contrasting treatment-phenotypes, SVA vs. CNA, we observed a significant increase in the abundance of 12 bacterial genera in SVA, including *Promicromonospora*, *C7867-002, Aequorivita*, *Nocardioides_A_392796*, and VFJQ01 (family level) (p < 0.05). The fungal genus *Penicillium* also exhibited increased abundance under these conditions.

On the other hand, four bacterial genera, *Stenotrophomonas_A_614440, Arachidicoccus, Azospirillum, Calidifontimicrobium*, showed reduced relative abundance in SVA, suggesting both water sensitivity and dependence on the host’s physiological status. Among fungi, *Cortinarius* and *Mortierella* also exhibited lower relative abundance in SVA.

## IV. Discussion

### Autumn drought as a distinct ecological regime

Our experimental design decoupled water deficit from thermal stress to address a structural bias in Mediterranean drought research. Ecophysiological studies on conifers such as *Pinus halepensis* have focused predominantly on summer, when drought coincides with high temperatures and reduced physiological activity, confounding heat and water stress in systems already operating under constrained carbon flux. By imposing progressive soil drying under moderate autumn temperatures (5–25 °C), we show that severe physiological decline can occur during an active growth phase: at 45% field capacity, stomatal conductance dropped to one-third of control values alongside near-complete growth cessation. Drought impact is therefore primarily governed by phenological state rather than heat (Forner et al., 2018; Zheng et al., 2026).

We propose a phenology-dependent drought framework in which drought during active growth constitutes a distinct ecological regime. *Pinus halepensis* exhibits a bimodal growth pattern with a secondary peak in autumn (de Luis et al., 2011; Valeriano et al., 2023), meaning that delayed or reduced autumn rainfall intercepts trees during a period of active carbon assimilation and rhizospheric carbon flux under these conditions. Root exudation under these conditions shapes microbial community assembly (Williams and De Vries, 2020; Zhalnina et al., 2018), meaning drought acts not only as a physiological constraint but as a selective force on microbiome assembly (Joseph et al., 2020; Oppenheimer-Shaanan et al., 2022a), making autumn drought a coupled plant–microbiome filtering event, likely more dynamic than drought occurring during summer quiescence.

This is directly relevant to reforestation: *P. halepensis* seedlings are typically outplanted in autumn, when establishment depends on timely rainfall onset (Taïbi et al., 2017). Under climate change, temporal variability in autumn precipitation (Vicente-Serrano et al., 2025) expose seedlings to drought during active growth, amplifying establishment failure risk. Phenology thus emerges as a primary axis of drought response, with concrete implications for predicting forest seedling establishment and forest regeneration under shifting Mediterranean climates.

The iCAMP analysis provides quantitative support for drought acting as an environmental filter on microbiome assembly. In the active fraction, reflecting metabolically engaged taxa, deterministic turnover was strongly associated with water regime: drought-treated plants showed nearly double the deterministic assembly of well-watered controls (35.5% in SNA vs. 19.6% in CNA), while drift was least important precisely in plants exposed to drought without visible physiological symptoms (SNA). In the resident fraction, deterministic processes increased progressively with symptom severity, from 29% in controls to 37.2% in severely affected plants, linking microbiome assembly mechanisms directly to host physiological state. Together, these patterns are consistent with drought acting as a selective force that shifts community assembly from stochastic toward deterministic processes, and suggest that this shift is mediated both by water availability and by the phenotypic condition of the host plant.

### Binary recovery and gas-exchange decoupling

Linking phenotype classes to gas-exchange traits clarifies why recovery was binary and *g_s_*–*Aₙₑₜ* became decoupled. Within the drought-treated cohort, a near-binary separation emerged between recovering and non-recovering individuals, a pattern obscured by mean recovery values. Critically, non-recovering plants were indistinguishable from future survivors under pre-stress conditions, showing comparable size, morphology, and initial gas exchange. This invisible susceptible subpopulation is ecologically significant: average physiological recovery metrics underestimate establishment failure risk, and the proportion of individuals lost to drought may be predictable neither from pre-planting traits nor from mean treatment responses.

Partial physiological recovery following rewatering is well documented in conifers. In, *Pinus halepensis* seedlings, g*_s_* remained 30–40% below pre-drought values after rewatering, while *A_net_* either fully recovered or remained reduced by 40% depending on the provenance. In *Pinus sylvestris* seedlings, both *g_s_* and *A_net_* recovered synchronously after drought, but remained at approximately 50% of control value, likely due to embolism-induced hydraulic limitations in stem (Rehschuh et al., 2020). Hydraulic recovery through embolism repair does not guarantee full photosynthetic recovery, as residual diffusional and biochemical limitations (i.e reduced mesophyll conductance and impaired Rubisco activity) can persist independently after rewatering (Flexas et al., 2004; Rehschuh et al., 2020)

Decoupling between *g_s_* and *A_net_* is also commonly observed under heat stress, where increased transpiration maintains leaf temperatures within safe limits (Gauthey et al., 2024), and can be amplified in drought-treated seedlings (Marchin et al., 2023). Because air temperatures remained within 5–25 °C throughout the experiment (Figure 2A), the g*_s_*–Aₙₑₜ decoupling observed here cannot be attributed to the lack of transpirational cooling as under heat stress (Gauthey et al., 2024). Instead, it likely reflects residual hydraulic constraints (embolism repair), biochemical limitations, and/or microbiome-mediated modulation of gas exchange as discussed below.

Gas exchange was measured on the apical shoot - the plant part showing the most severe visual symptoms, while lateral shoots showed less clear phenological response, likely experienced less intense drought stress, potentially maintaining better physiological functions and sustained carbohydrate fluxes to the rhizosphere. The binary recovery pattern, in which only apical shoots fail to recover is not well documented. One mechanistic hypothesis is relate to apical dominance. Damage to the apical dominance may disrupt hormonal balance (Beveridge et al., 2023), potentially altering carbon allocation and root development. Supporting this view, oak seedlings grown under shelter developed stronger apical dominance and invested less in root development, whereas unsheltered seedlings allocated more resources to broader root systems (Mariotti et al., 2015). Drought-induced disruption of apical dominance may redirect belowground carbon allocation and rhizodeposition, with downstream consequences for root-associated microbiome assembly.

### Microbiome-mediated modulation of gas exchange

Leaf gas exchange can also be modulated by root-associated microbiomes, including ectomycorrhizal fungi and plant growth-promoting bacteria, under both stressed and unstressed conditions. Our differential abundance analysis reveals patterns consistent with this. The ectomycorrhizal genus *Cortinarius*, whose symbiotic associations with Pinus roots support host nitrogen and water economy (Heinonsalo et al., 2015), was enriched in drought-exposed plants that maintained physiological function (SNA) but depleted in severely affected plants (SVA), mirroring the broader decline in symbiotrophic fungi across drought-induced phenotypes. Conversely, saprotrophic taxa including *Mortierella* were enriched under drought, consistent with community-level increases in saprotrophic fungal guilds, and *Mortierella* in particular has been associated with plant stress tolerance (Ozimek and Hanaka, 2020). In SNA plants, the co-enrichment of *Pseudomonas and Trichoderma*, commonly considered as plant growth-promoting microorganisms capable of modulating stomatal aperture and (Contreras-Cornejo et al., 2009; Vurukonda et al., 2016), may further explain active microbiome-mediated support of gas exchange in drought-resistant individuals. At the community level, functional predictions indicate elevated ureolytic and nitrogen-fixing bacterial activity in the active fraction of drought-treated plants, suggesting a reorganization toward nutrient mobilization under water deficit, though the specific taxa driving this signal require further resolution. Together, these patterns suggest that the microbiome contribution to gas exchange maintenance under drought is phenotype-dependent, with resistant plants recruiting a functionally distinct microbial community.

### Drought increases root microbiome diversity in Pinus halepensis

Having established that drought severity and recovery capacity diverge across phenotype classes, we next examined whether these physiological trajectories were reflected in the structure and diversity of root-associated microbial communities.

Contrary to expectations, drought-submitted *P. halepensis* plants showed significantly higher bacterial richness and evenness, compared to well-watered control plants. Drought is typically associated with reduced microbial diversity due to habitat filtering and resource limitation (Kristy et al., 2022; Yang et al., 2021; Zhou et al., 2020), making this result unusual. We hypothesize, that increased root exudation under drought enriches the rhizosphere with low-molecular-weight compounds that serve as substrates for microbial growth, thereby promoting microbial proliferation and diversity (Oppenheimer-Shaanan et al., 2022b; Preece et al., 2018; Tiziani et al., 2022; Ulrich et al., 2022). Cases of drought-induced alpha diversity increase are rare, and have been reported under soil drought legacy effects in *Olea europea* (Preece et al. 2019), *Salix purpurea* and drought-sensitive bean cultivars (Silva et al., 2025). Evidence from tree systems further suggests that drought can reshape root microbiome composition without reducing richness (Addison et al., 2025), consistent with a process of community reorganization rather than drastic diversity loss — a pattern our results extend and amplify by showing an active fraction richness increase under autumn drought in an active growth phase. Whether this diversity increase reflects genuine recruitment of new taxa or the release of previously suppressed low-abundance members under altered resource regimes remains to be determined, but it points unambiguously to drought as a driver of microbiome expansion rather than contraction in this system.

### Drought induces Pinus halepensis’ microbiome restructuring in phenotype-related manner

Microbiome-phenotype associations have rarely been explored as a framework linking the functional profiles of both host and microbiome to their respective compositional and phenotypic traits. However, the emerging concept of microbiome-assisted breeding highlights the potential of harnessing such associations by integrating desired plant phenotypes with supportive microbiome compositions (Cernava, 2024; Mueller and Linksvayer, 2022).

In the present study, we observed a clear entanglement between *P. halepensis* drought phenotype and associated microbial community in both active and resident microbial fraction, each representing distinct functional states. In the resident bacterial fraction, significant differences were detected in both weighted and unweighted UniFrac metrics, suggesting that phenotype-associated community shifts involved not only changes in taxonomic composition (presence/absence) but also alterations in relative abundance. This indicates a broad restructuring of the microbial community in response to drought-induced plant phenotypes. By contrast, in the active fraction, only the weighted Unifrac distance was significant, implying that the observed phenotype-microbiome linkage was primarily driven by shifts in the relative abundance of established taxa, rather than species turnover. These trends are corroborated by MiRKAT-MC: in resident fraction, both phylogenetic and non-phylogenetic metrics were significant, whereas in active fraction, non-phylogenetic distances dominated, with only weighted UniFrac remaining significant—again pointing to abundance-driven shifts in the active fraction.

This pattern is consistent with the PERMANOVA results, where the interaction between water regime and phenotype explained the largest share of variance in both resident (29.8%) and active (22.0%) fractions, whereas the phenotype alone was not significant (Table 1).

To our knowledge, this is the first report of a direct, phenotype-related restructuring of both resident and active root-associated bacterial communities in a conifer. While most plant–microbiome links are investigated via GWAS (Deng et al., 2022), recent work in *Medicago truncatula* showed that rhizosphere bacterial variation is closely associated with ecophysiological and genetic plant traits (Zancarini et al., 2025). In human systems, microbiome–phenotype association studies are far more advanced (Awany et al., 2019; Kurilshikov et al., 2021).

Because seedlings originated from a mixed seed lot, we cannot fully disentangle drought-induced phenotypic divergence from underlying genetic or epigenetic differences; such host variation could contribute to the observed phenotype–microbiome coupling via constitutive differences in exudation or immune signalling. Future work using clonal material or genotyped half-sib families will be necessary to partition genetic/epigenetic vs. environmental contributions to microbiome assembly.

### Contrasting assembly mechanisms link drought severity to functional reorganization of the root microbiome

The iCAMP analysis revealed contrasting assembly trajectories for bacterial and fungal communities respond to drought through distinct assembly mechanisms, highlighting fundamental differences in how these two kingdoms respond to drought stress. In the bacterial community, drought promoted increasingly deterministic assembly as phenotype severity increased, with deterministic processes rising from ∼29–30.5% in non-affected plants to 37.2% in severely affected plants. This indicates that environmental filtering strengthens with host physiological impairment, favouring stress-adapted bacterial taxa through predictable selection processes. Drought, disrupts nutrient flows and root-soil interactions, likely creating a selective pressure on the bacteria community that is well documented in plant associated microbiomes (Li et al., 2023; Sun et al., 2025), and typically interpreted as stronger environmental filtering (Moore et al., 2023; Wang et al., 2023).

We propose two non-exclusive mechanisms driving this shift. First, declining water potential and altered soil structure create increasingly selective abiotic conditions that filter drought-tolerant taxa (Hayashi et al., 2024; Stegen et al., 2012). Second, enhanced root exudation under drought may provide an additional host-mediated selection force, as plants release specific metabolites that selectively promote beneficial microorganisms capable of supporting stress tolerance (Sasse et al., 2018; Zhalnina et al., 2018). It should be noted that of the ‘deterministic’ bacterial filtering we detected could also reflect constitutive differences among host genotypes rather than drought effects per se, a limitation discussed above.

The divergence in assembly modes mirrors the diversity patterns described above: deterministic filtering coincided with increased in bacterial α-diversity, suggesting that stronger environmental selection does not necessarily reduce richness but instead reorganizes community membership around a more diverse but functionally constrained set of taxa.

Fungal communities exhibited the opposite response, shifting toward stochastic assembly mechanisms under drought stress, especially in the active fraction. The sharp increase in drift processes from 43.6% to 72.3% in severely affected plants suggests that drought conditions disrupt fungal community organization, likely through fragmentation of hyphal networks and reduced connectivity. Interestingly, a recent paper by (Pan et al., 2024) indicate that drought increased the part of deterministic process; however, they found an increase in stochasticity in the bulk soil. We hypothesize, that our sampling captured an immediate disturbance in which fungal communities were actively reorganizing, and that stabilization toward more deterministic assembly would occur over a longer recovery period

As drought advances, reduced photosynthesis and declining carbon allocation to roots alter the quality and quantity of rhizodeposits available to microbial communities (Butler et al., 2003; Pausch and Kuzyakov, 2018). Under prolonged stress, depletion of labile carbon substrates and progressive degradation of root tissues likely shift community composition toward taxa capable of utilizing more recalcitrant structural polymers (Amundrud et al., 2019). This shift is reflected in the functional guild data: the proportion of saprotrophic fungi such as wood and soil saprotrophs increased across drought-induced phenotypes, while symbiotrophic fungi declined, consistent with disruption of mutualistic associations and opportunistic exploitation of degrading root tissues. The stochastic assembly observed in fungi is therefore not simply random — it reflects a functional reorganization driven by the collapse of host-mediated structuring and the rise of resource-driven opportunism.

These compositional and assembly shifts are further supported by FAPROTAX functional predictions. Higher proportions of ureolytic and nitrogen-fixing bacteria were observed in drought-induced phenotypes (SNA, SMA, SVA) compared to controls, particularly in the active fraction, while nitrate reduction and respiration peaked in CNA in the resident community (Figure 6A). This functional enrichment suggests that deterministically assembled bacterial consortia under drought emphasize rapid nitrogen turnover, potentially compensating for reduced nitrogen availability under limited carbon flux and altered root physiology. The concurrent rise of ureolysis and N-fixing groups under drought (Figure 6A) supports this deterministic filtering, as resource scarcity and altered rhizodeposition likely favour bacteria with specific nutrient-cycling capacities (Naylor et al., 2022; Santos-Medellín et al., 2017). In contrast, fungal communities shifted toward stochasticity while saprotrophic guilds increased and symbiotrophs declined (Figure 6B), consistent with hyphal disruption and carbon-starved roots shedding tissues that opportunistic decomposers can exploit (De Vries et al., 2020; Hawkes and Keitt, 2015). Together, these bacterial and fungal patterns point to a coherent but asymmetric restructuring of the root microbiome under drought: bacteria becoming more predictably selected and functionally specialized around nutrient cycling, while fungi become more compositionally variable and opportunistically saprotrophic as drought severity increases.

### Water Deficit reshapes pine root microbiome network topology

Water deficit fundamentally altered the architecture of root microbiome co-occurrence networks, shifting community organization from redundancy to fragmentation. Under well-watered conditions, networks exhibited greater complexity but lower density, indicative of diverse yet loosely connected microbial interactions. High complexity of this kind can enhance resilience to environmental perturbations through functional redundancy, distributing resources broadly across the network (Mougi and Kondoh, 2012). Under drought, networks became smaller and denser, but also more modular and fragmented (particularly in SNA), indicating communities that are internally cohesive yet poorly connected across modules (Figure 5). Such compartmentalization can buffer local disturbances, but it also limits cross-module support, making the whole system easier to tip if several modules fail simultaneously (Banerjee et al., 2018; De Vries and Shade, 2013; Gilarranz et al., 2017; Pilosof et al., 2017). Drought did not simply shrink the networks; it changed how information and resources can move through them.

The most ecologically significant topological shift was the loss of *Rhizobium* as the keystone hub taxon in drought-affected phenotypes. Under well-watered plants, *Rhizobium* occupied a central position, orchestrating community interactions, with *Pararhizobium* acting as an important connector (Supplementary Figure 6). Both taxa contribute to nitrogen cycling through their fixation capabilities (Tanunchai et al., 2023), and Rhizobium plays established roles in plant growth promotion the production of signalling molecules that shape both microbial dynamics and plant physiological responses (Lindström and Mousavi, 2019). The loss of *Rhizobium* as a keystone hub in drought phenotypes is ecologically meaningful. The displacement of this mutualistic hub forces the network to reorganize around less influential connectors, e.g. *Sphingobium*, *Pararhizobium*, *Achromobacter*) - taxa associated with degradation of complex organic compounds rather than mutualistic plant support (Banerjee et al., 2018; Toju et al., 2018). This topological shift is coherent with the functional predictions (FAPROTAX/FUNGuild): nitrogen -cycling bacteria persisted but redistributed across the network, while saprotrophic fungi rose while symbiotrophs declined. The network no longer hinged on one mutualistic hub; it pivoted to opportunistic decomposers and smaller bacterial connectors, matching the carbon-and nitrogen-use strategies expected under water deficit (Santos-Medellín et al., 2017).

Across phenotypes, increasing drought severity (SMA → SVA) was mirrored by progressive network contraction and centralization: betweenness and eigenvector centralities concentrated in fewer taxa, suggesting that deterministic filtering at the taxon level translated into topological “filtering” of the network itself. Notably, SNA plants without visible symptoms already showed high modularity and low robustness, implying that microbiome networks respond before plant physiology collapses (early-warning signal). This early-warning signal is consistent with the iCAMP results showing that deterministic assembly was already elevated in SNA relative to CNA.

Random-removal robustness ranking (CNA > SMA > SVA > SNA) indicated that drought networks, especially SNA, are fragile to stochastic taxon loss which is consistent with their smaller, tightly connected topology. Paradoxically, targeted-removal vulnerability was lowest in SNA, meaning once big hubs are gone (as in SNA), there is less to target. Functionally, that is a double-edged sword or trade-off: fewer critical hubs reduce catastrophic collapse risk but also reduce the system’s capacity to re-route functions when individual nodes are lost.

Fungal taxa were consistently peripheral across all phenotypes, with only one unidentified Ascomycota in SMA occupying a connector position. This peripheral topography matches the observed guild shifts: under drought, fungi shifted from mutualistic symbiotrophs to opportunistic saprotrophs.

Taken together, the taxonomic, functional, and topological network changes described a microbiome-driven adaptation of *Pinus halepensis* seedlings to autumn drought. The microbiome of drought-resistant plants (SNA) was characterized by higher diversity, more deterministic bacterial assembly, and network reorganization that preceded visible physiological stress — consistent with an early, active microbial response to water deficit rather than a passive consequence of host decline. The fact that these changes differentiated active from resident fractions reinforces that metabolically engaged communities responded first and most strongly to drought, underscoring the high reactivity of the root-associated microbiome to soil water availability and host physiological state.

## V. Conclusions

Our results provide a multifaceted view of drought impacts at moderate temperatures in *P. halepensis* seedlings, with implications for both plant physiology and microbiome ecology. Autumn drought triggered rapid physiological responses and induced a functional decoupling between stomatal conductance and CO_2_ assimilation that became fully apparent during recovery, a pattern distinct from heat-driven decoupling and attributable to residual hydraulic and biochemical limitations.

At the microbiome level, distinct drought-induced phenotypes were associated with specific microbial assemblages, shaped through contrasting mechanisms in metabolically active vs. resident communities. Bacterial communities were predominantly governed by deterministic processes, as host physiological impairment intensified, reflecting progressive environmental and host-mediated filtering toward stress-tolerant taxa. Fungal communities followed the opposite trajectory, shifting toward stochastic drift as symbiotrophs declined and saprotrophic guilds expanded. At the network level, drought destabilized cross-kingdom interaction architecture, displacing *Rhizobium* from its central hub position and forcing reorganization to the smaller, less influential connectors. This topological shift that preceded visible plant stress in drought-resistant individuals and may represent an early warning signal of community fragility.

Together, these findings demonstrate that even short, severe drought at moderate temperatures, can decouple host physiology and restructure root microbiomes through deterministic bacterial filtering, stochastic fungal drift, and rapid network rewiring. The shift from mutualistic to resource-scavenging microbial strategies under water deficit is an apparent strategy of microbiome adaptation to drought. These mechanisms should be considered when preparing seedlings for Mediterranean reforestation, especially as climate change increases the frequency and unpredictability of autumn droughts that threaten seedling survival.

While our design captured realistic genetic heterogeneity of nursery stock and growth practises, it also limits our ability to separate host genetic and epigenetic effects from drought-driven filtering. Future work using clonal material or genotyped half-sib families will be necessary to disentangle host physiological and microbiome-driven adaptation.

## Author Contributions

Conceptualization, C.S.; Methodology, C.S., J.R., M.d.C., I.M.R. and I.A.; Software, I.A.; Validation, C.S., and I.M.R.; Formal Analysis, I.A.; I.M.R. and C.S.; Investigation, I.A., and I.M.R.; Resources, C.S., I.M.R., J.R., M.d.C.,; Data Curation, I.A. and I.M.R.; Writing—Original Draft Preparation, I.A.; Writing—Review and Editing, C.S., I.M.R.; Visualization, I.A., I.M.R. and C.S.; Supervision, C.S.; Project Administration, C.S.; Funding Acquisition, C.S. and I.M.R. All authors have read and agreed to the published version of the manuscript.

## Acknowledgments and funding

This research was funded through the RESTORE project, 2019–2020 BiodivERsA+ joint call for research proposals under the BiodivClim ERA-Net COFUND program and with the following funding organisations: Fundação Araucária/Secretaria de Estado da Ciência, Tecnologia e Ensino Superior do Paraná (NAPI Biodiversidade), FAPESP (Brazil), French National Research Agency (ANR) (grant number ANR-20-EBI5-0008-07), Federal Ministry of Education and Research (Germany).

I.A. received additional funding from the HORIZON EUROPE INFRA-2022-TECH project ‘PHENET’, grant number 101094587. This work was supported by the CAPES-COFECUB program (project N° Sv 946/10-41880PL), which facilitated international collaboration and researcher mobility. Special thanks to the PNRGF-ONF (Office National des Forêts) for their expertise and for providing the facilities to grow our tree seedlings. Huge thanks to Bastian Charillat, for the care and attention to the tree seedlings during their development.

