## Supplementary figure 1 for "Autumn drought drives deterministic bacterial filtering and network destabilization in a phenotype-related manner in *Pinus halepensis* seedlings"

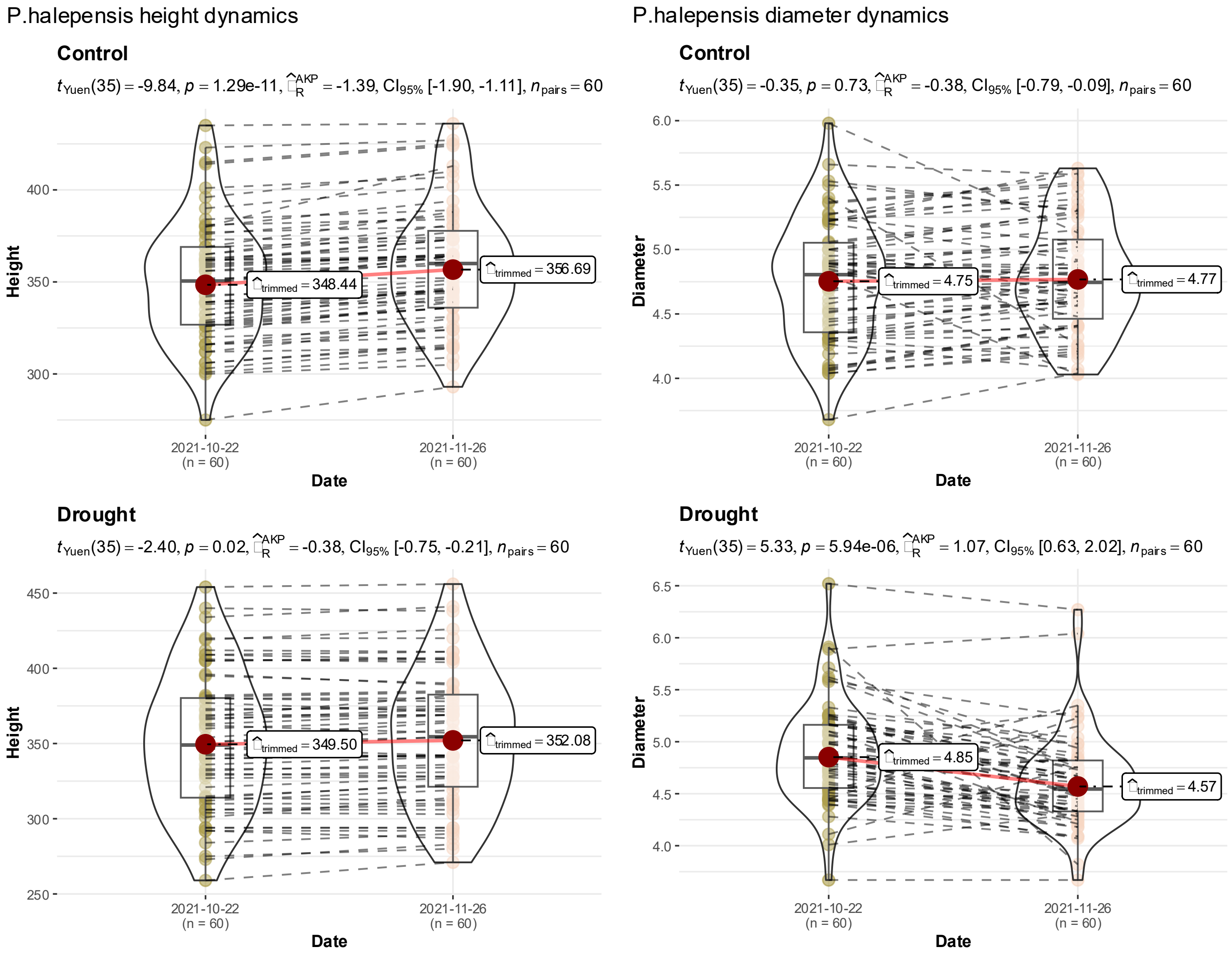


**Supplementary figure 1.** Height and diameter changes between beginning and end of experiment. Drought impact on height and diameter increase was tested using paired Yuen test for trimmed means at $p < 0.05$.
