## Supplementary figure 2 for "Autumn drought drives deterministic bacterial filtering and network destabilization in a phenotype-related manner in *Pinus halepensis* seedlings"

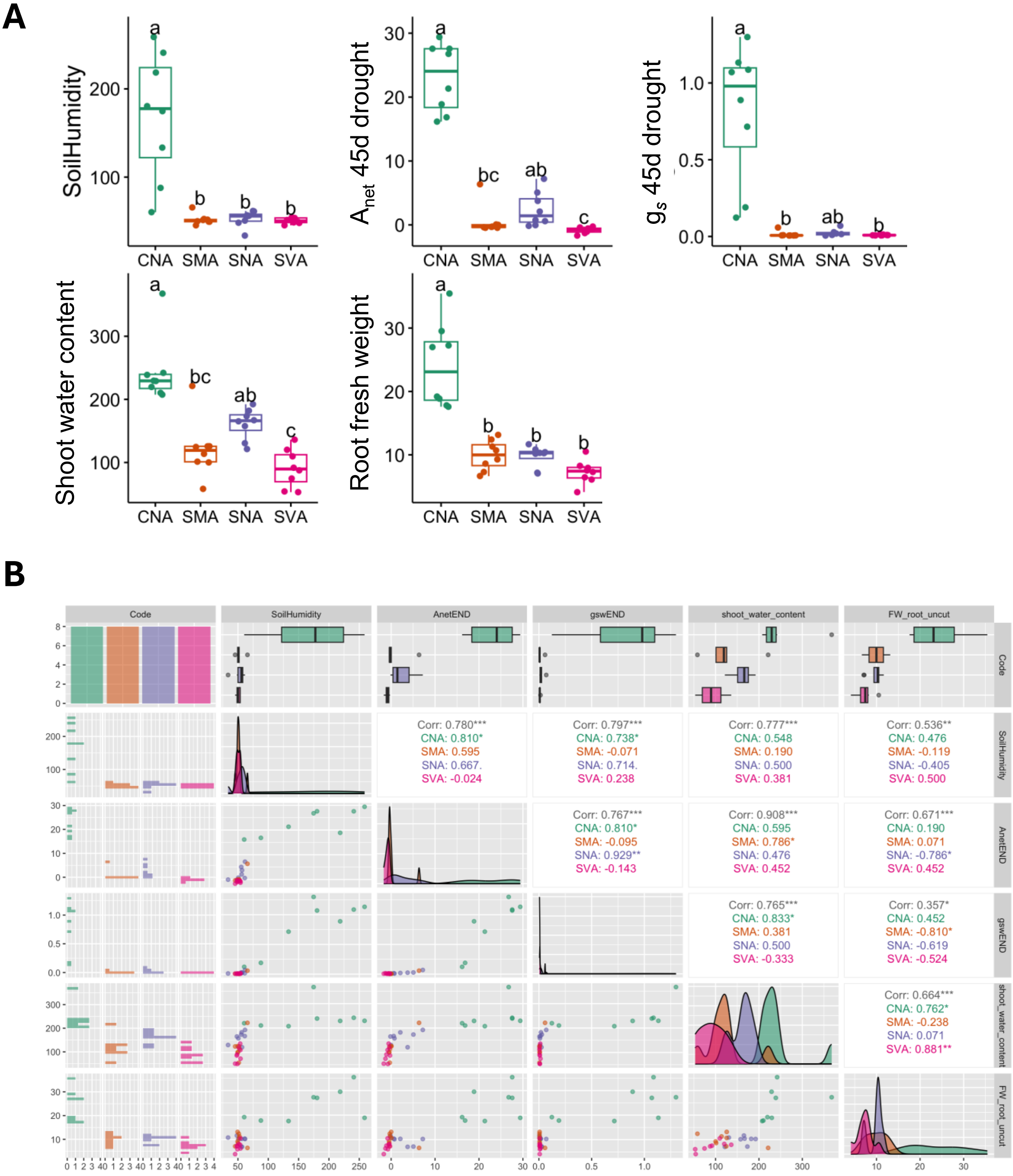


**Supplementary figure 2.** Measured physiological and environmental parameters. (A) Box plot showing the differences between treatment-phenotypes. (B) Variables Spearman correlation plot. Letters represent Kruskal-Wallis test result with Dunn multiple comparisons test at $p < 0.05$.
