## Supplementary figure 3 for "Autumn drought drives deterministic bacterial filtering and network destabilization in a phenotype-related manner in *Pinus halepensis* seedlings"

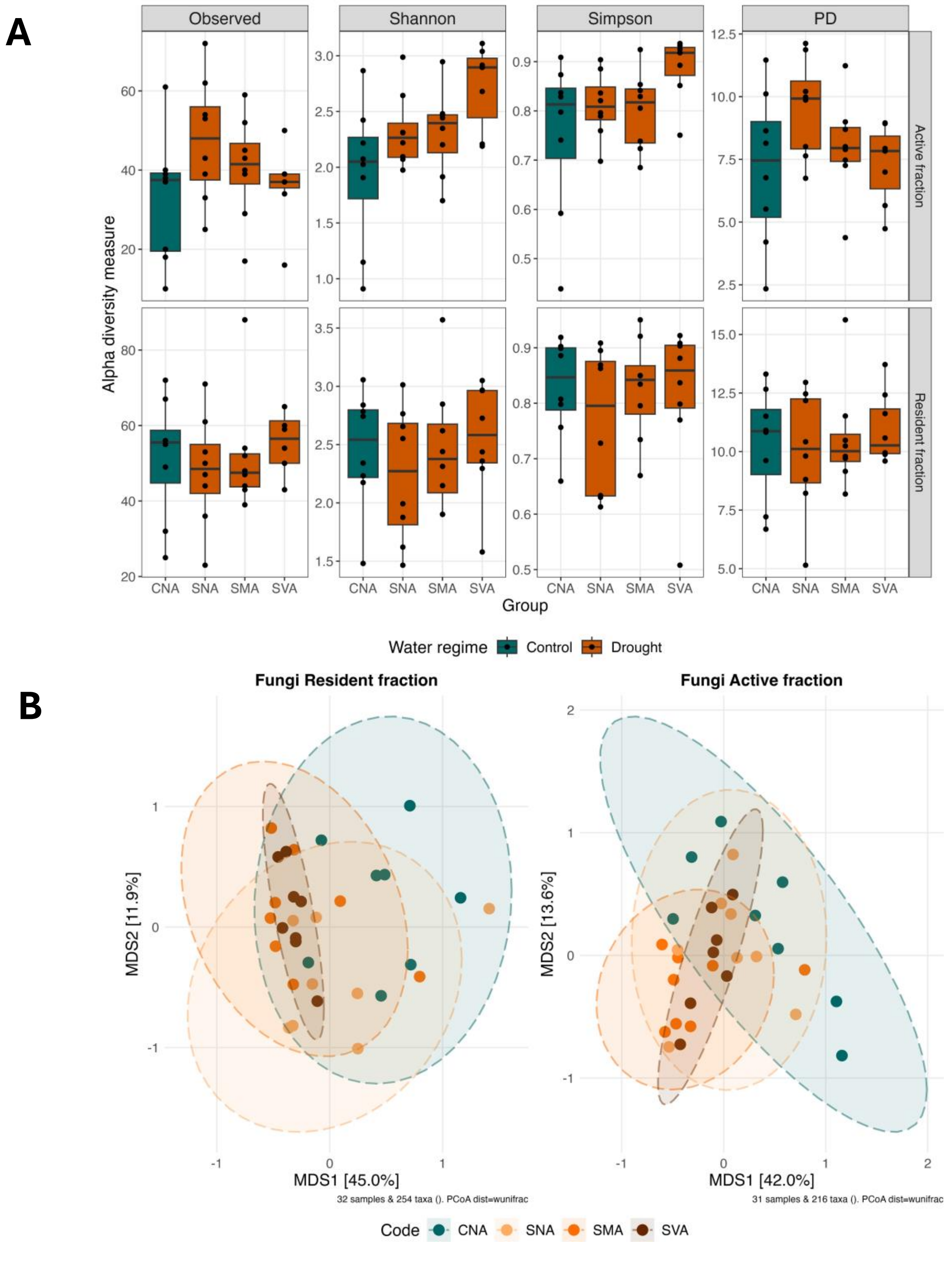


**Supplementary figure 3. Fungal diversity analysis of associated with *Pinus halepensis* in different treatment-phenotypes**. **(A)** Alpha diversity metrics. **(B)** Beta diversity analysis using PCoA of weighted UniFrac distances for active and resident microbial fractions. No significant differences were detected in alpha diversity metrics.
