## Supplementary figure 4 for "Autumn drought drives deterministic bacterial filtering and network destabilization in a phenotype-related manner in *Pinus halepensis* seedlings"

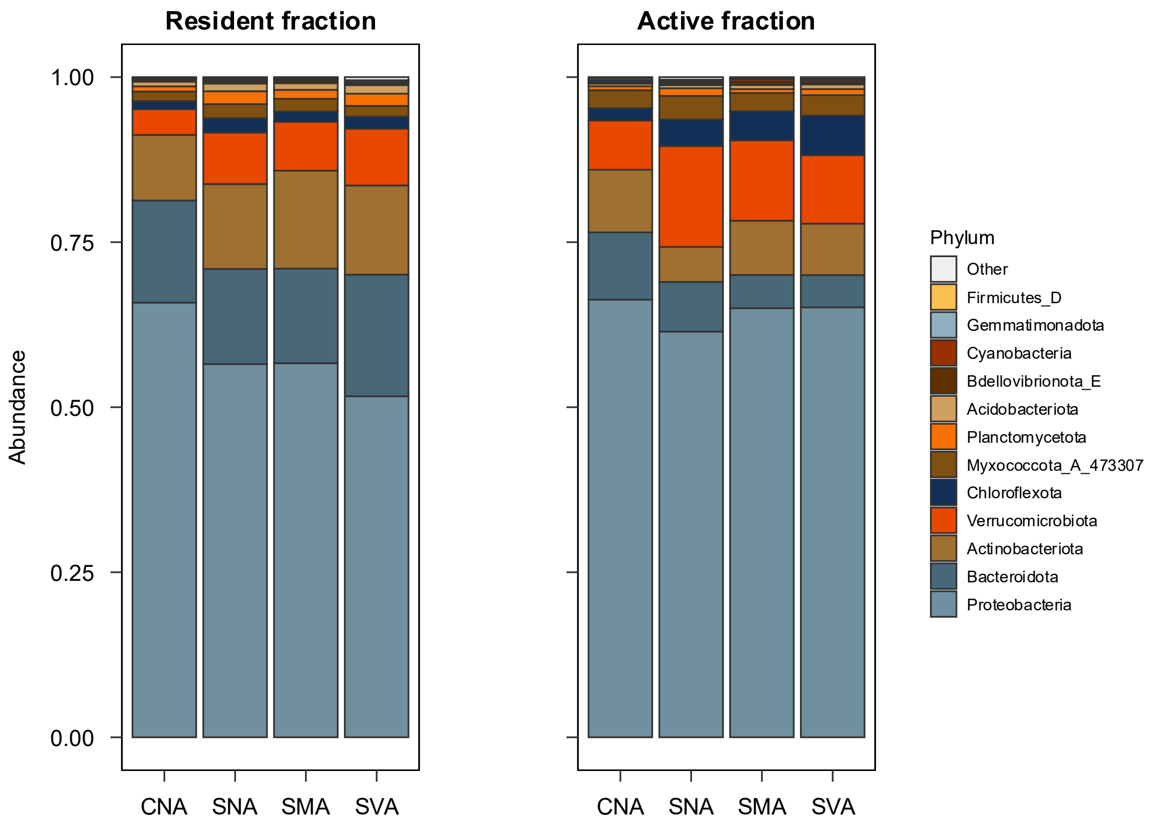


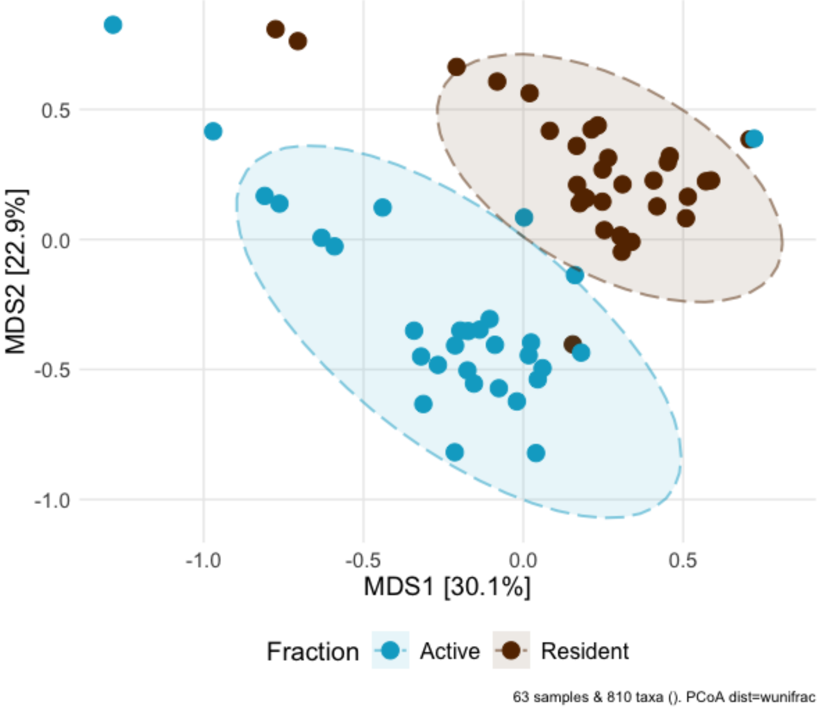


**Supplementary figure 4 Taxonomic composition and community dissimilarities between resident and active microbial fraction of Pinus halepensis roots.** (A) Barplot for taxonomic composition of bacteria at Phylum level in different treatment-phenotypes (B)
