## Supplementary figure 5 for "Autumn drought drives deterministic bacterial filtering and network destabilization in a phenotype-related manner in *Pinus halepensis* seedlings"

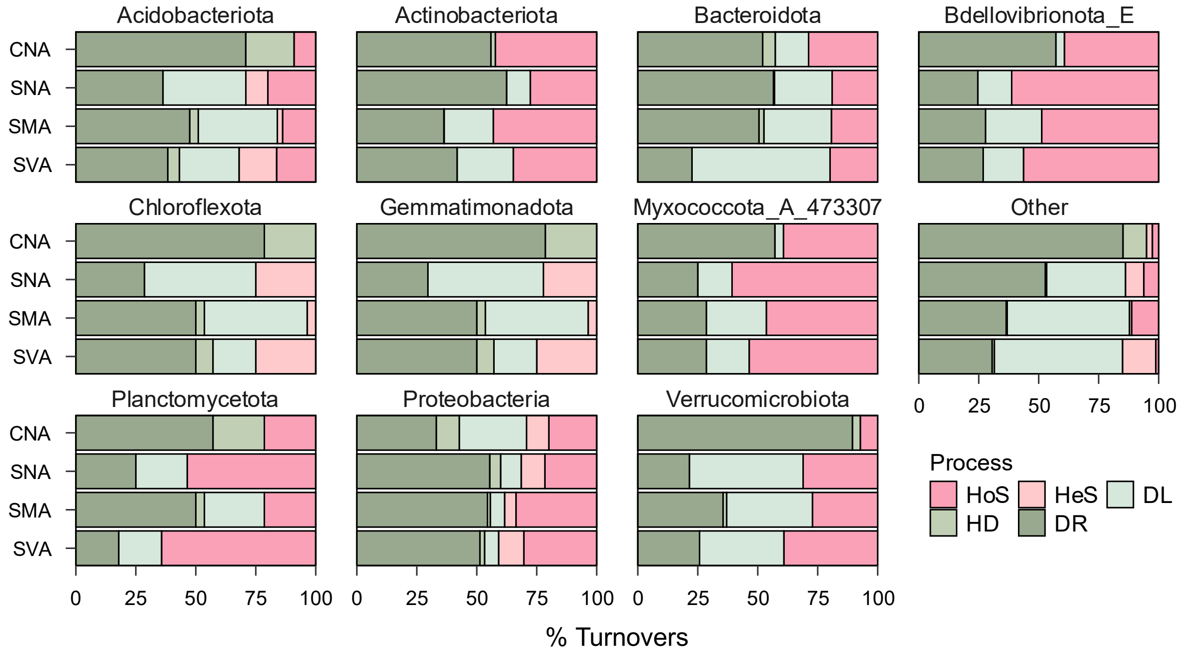


**Supplementary Figure 5** Ecological assembly mechanisms of resident (gDNA-based) microbial communities associated with *Pinus halepensis* treatment-phenotypes. Relative contributions of community assembly processes within each treatment-phenotype, belonging to homogeneous selection (HoS), heterogeneous selection (HeS), dispersal limitation (DL) homogenizing dispersal (HD), or drift (DR) the top ten the most abundant microbial taxa.
