## Supplementary figure 6 for "Autumn drought drives deterministic bacterial filtering and network destabilization in a phenotype-related manner in *Pinus halepensis* seedlings"

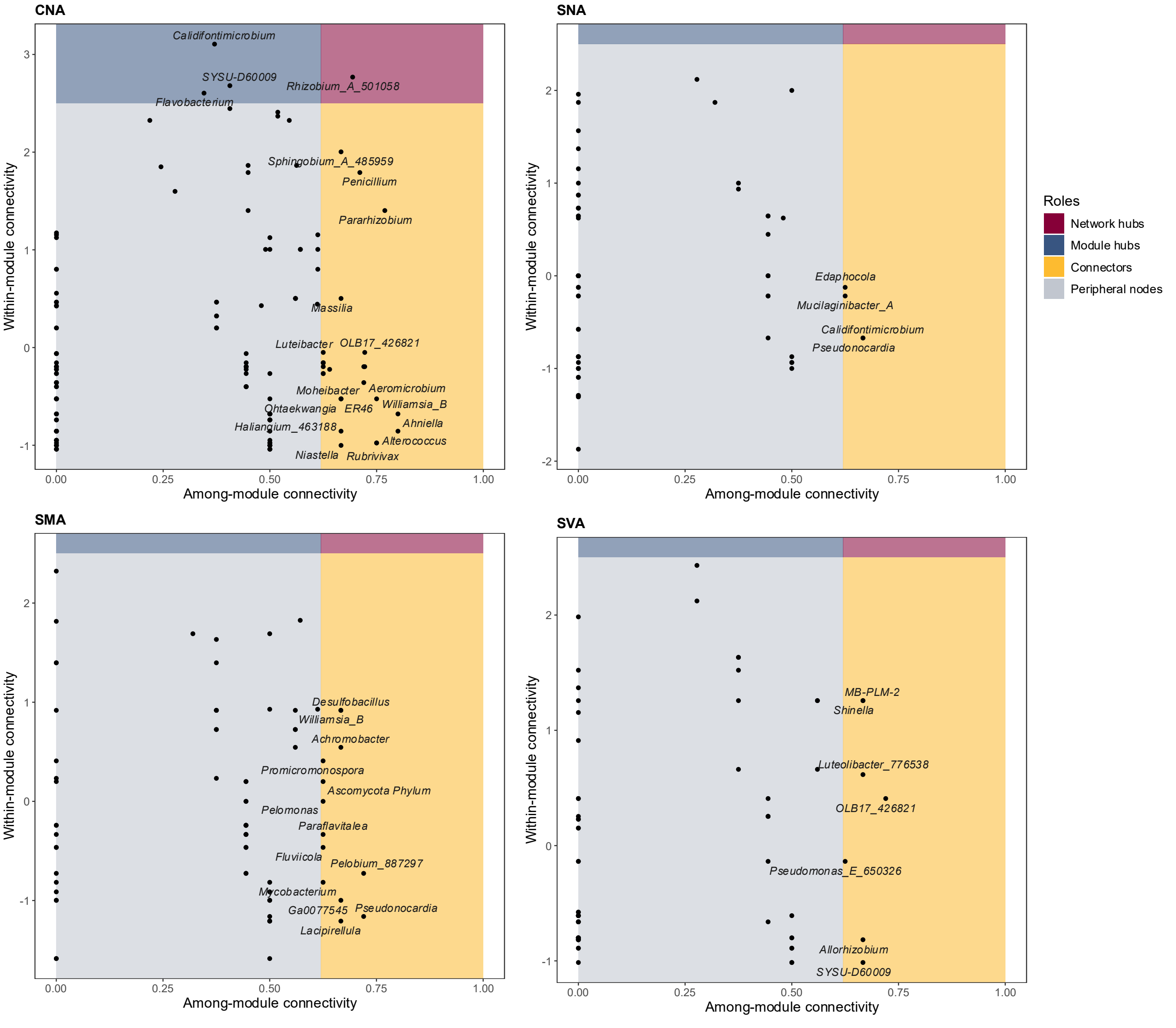


**Supplementary Figure 6. Zi-Pi plot showing the distribution of taxa based on their topological roles.** Zi-Pi plots which consisted of axes describing within module connectivity (Zi) and among module connectivity (Pi) metrics. Nodes are classified based on their Zi and Pi scores (2.5 and 0.62 respectively) into four ecological roles: peripheral nodes, or specialists (less influential taxa; grey panel), module hubs (represent taxa with high influence locally; blue panel), connectors (represent taxa important for cross-module interactions; yellow panel) and network hubs (identifying keystone taxa that maintain microbial community stability; red panel).
