## Supplementary figure 7 for "Autumn drought drives deterministic bacterial filtering and network destabilization in a phenotype-related manner in *Pinus halepensis* seedlings"

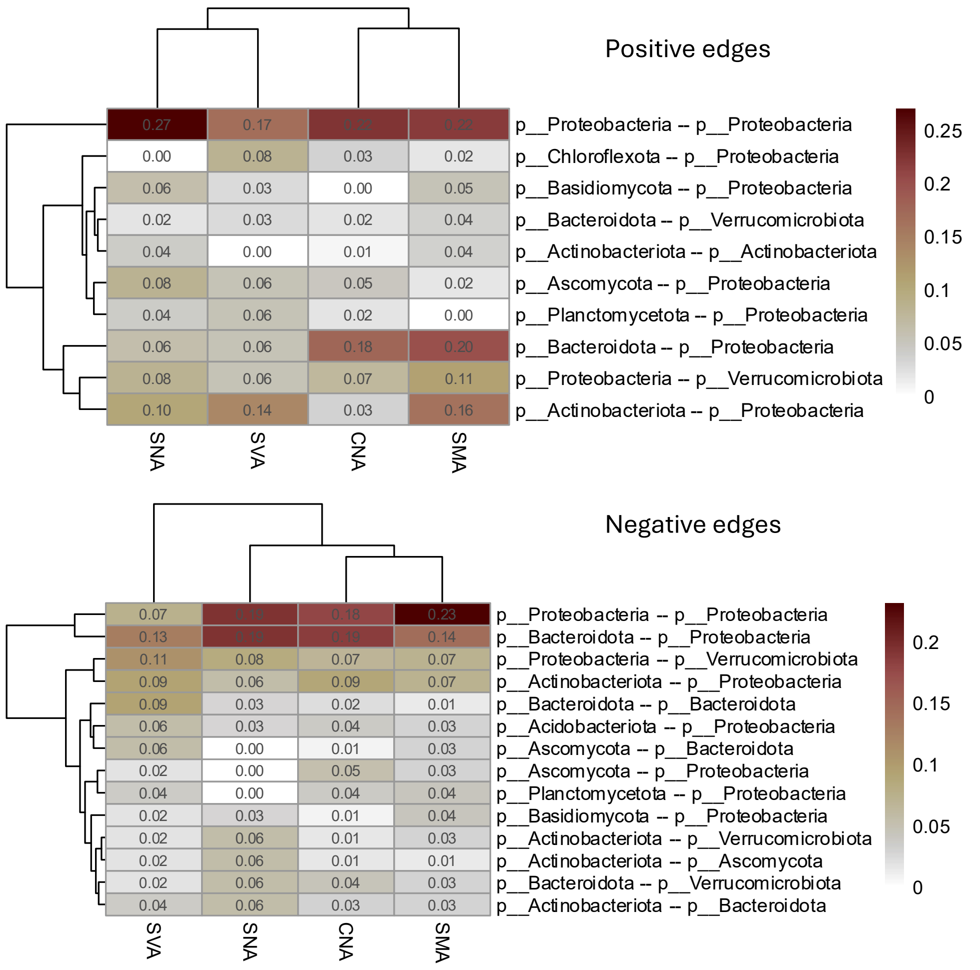


**Supplementary Figure 7.** Comparison of node sources of microbial co-occurrence edges across networks in *Pinus halepensis*. The figure distinguishes whether each edge connects taxa from the same phylum or from different phyla. Positive associations (top panel) and negative associations (bottom panel) are shown separately. Edges were filtered to retain only those with strength > 0.02. Proportions were normalized by the total number of positive or negative edges per network to allow cross-network comparison.
